# A Second Drug Binding Site in P2X3

**DOI:** 10.1101/2024.06.10.598171

**Authors:** Trung Thach, KanagaVijayan Dhanabalan, Prajwal Prabhakarrao Nandekar, Seth Stauffer, Iring Heisler, Sarah Alvarado, Jonathan Snyder, Ramaswamy Subramanian

## Abstract

Purinergic P2X3 receptors form trimeric cation-gated channels, which are activated by extracellular ATP. P2X3 plays a crucial role in chronic cough and affects over 10% of the population. Despite considerable efforts to develop drugs targeting P2X3, the highly conserved structure within the P2X receptor family presents obstacles for achieving selectivity. Camlipixant, a potent and selective P2X3 antagonist, is currently in phase III clinical trials. However, the mechanisms underlying receptor desensitization, ion permeation, principles governing antagonism, and the structure of P2X3 when bound to camlipixant remain elusive. In this study, we established a stable cell line expressing homotrimeric P2X3 and utilized a peptide scaffold to purify the complex and determine its structure using cryo-electron microscopy (cryo-EM). P2X3 binds to camlipixant at a previously unidentified drug-binding site and functions as an allosteric inhibitor. Structure-activity studies combined with modeling and simulations have shed light on the mechanisms underlying the selective targeting and inhibition of P2X3 by camlipixant, distinguishing it from other members of the P2X receptor family.

## Introduction

Under both physiological and pathophysiological conditions, various cell types release ATP (Corriden & Insel, 2010). In the lungs, extracellular ATP activates nerve terminals, which are pivotal for triggering the cough reflex (Zhang *et al*, 2022; Canning *et al*, 2014). Cough serves as a vital defense mechanism for the airways and protects against the inhalation or aspiration of harmful environmental particles. When a cough persists for eight weeks or longer without evidence of lung disease on radiographic imaging, it is classified as chronic cough (Bolser, 2006). This condition affects more than 10% of the population, with a notable proportion of individuals experiencing idiopathic, unexplained, or refractory cough, lacking a clear underlying cause (Kamei *et al*, 2005; Canning *et al*, 2014; Chung, 2011).

Neuronal signaling that prompts coughing is well documented and is primarily mediated by ligand-gated ion channels known as P2X3 receptors (Spinaci *et al*, 2021; Khakh & Alan North, 2006). These receptors, comprising an extracellular ligand-binding domain and a membrane-spanning ion channel domain, swiftly convert extracellular chemical signals into transmembrane ionic currents, facilitating rapid intercellular signaling and chemosensation (Sheng & Hattori, 2022; Finger *et al*, 2005). Purinergic P2X3 receptors gated by ATP exist in both homotrimeric (comprising three P2X3 subunits) and heterotrimeric (comprising one P2X2 subunit and two P2X3 subunits, termed P2X2/3) configurations. Although P2X3 has been implicated in pathological cough, P2X2/3 also plays a role in taste transmission (Spinaci *et al*, 2021; Ford & Undem, 2013). Antagonists targeting P2X3 hold promise for chronic cough treatment by attenuating neuronal hypersensitivity. However, designing drugs that selectively target P2X3 over P2X2/3 or other P2X receptors is challenging owing to the highly conserved structures of these receptors. For example, gefapixant, which acts as a negative allosteric modulator of P2X3 receptors, has shown encouraging outcomes in clinical trials targeting refractory chronic cough. Nonetheless, gefapixants can induce side effects, notably taste alteration or loss (Abdulqawi *et al*, 2015). This phenomenon arises from the dual action of gefapixant, which affects not only P2X3 homotrimers associated with refractory chronic cough, but also P2X2/P2X3 heterotrimers involved in taste signal transmission (Wang *et al*, 2018; Abdulqawi *et al*, 2015).

Recently, camlipixant has emerged as a potent and selective P2X3 antagonist, with significantly higher potency against P2X3 than against P2X2/3 (25 nM versus 24,000 nM potency in human cell lines). Camlipixant shows potential for cough relief with a minimal impact on taste perception and is currently undergoing phase III clinical trials (NCT05599191). Despite these promising results, the structural basis underlying the selective targeting of P2X3 by camlipixant is not yet understood. Although the crystal structures of P2X3 have elucidated common mechanisms, such as ATP and antagonist binding and gating within the P2X receptor family (Mansoor *et al*, 2016; Kasuya *et al*, 2017; Shen *et al*, 2023; Jiang *et al*, 2021) questions remain regarding the mechanism that makes camlipixant specific to P2X3. For instance, what are the structural determinants of camlipixant’s preference for P2X3 over P2X2/3?

Considering the importance of P2X receptors as drug targets, we identified and mapped the binding site of camlipixant on P2X3 using single-particle cryo-electron microscopy (cryo-EM). Our results shed light on the structural basis of the inhibitory action of camlipixant on P2X3. Our findings, supported by biochemical and biophysical experiments and Molecular Dynamics (MD) simulations, revealed a cryptic drug-binding site for camlipixant that interferes with the conformational changes associated with P2X3 receptor inhibition. The camlipixant-bound P2X3 complex structure provides a foundation for future drug discovery efforts targeting the P2X receptors.

## Results

### GnTI^-^ stable cell line expression of P2X3

Because of the challenges posed by high glycosylation and low expression levels of the P2X3 receptor in human cell membranes, we first expressed the P2X3 receptor in the Expi293F GnTI^-^ (N-acetylglucosaminyltransferase I-negative) suspension cell line via transient transfection using a plasmid containing the gene encoding P2X3 from *Canis lupus* sp. (cP2X3). *Canis lupus* P2X3 is more than 97% identical to human P2X3, and all key residues involved in ATP binding, drug binding, and the channel are conserved. This platform aims to enhance the yield of P2X3 by reducing the number of sugars added by glycosylation machinery. This expressed protein was more conducive to structural studies. The P2X3 construct displayed robust expression and the expressed protein was localized to the cellular membrane (Figure S1A). Subsequently, we generated a stable cell line expressing GFP-P2X3 using the PiggyBac transposon system (Figure S1B). The viability assay and GFP fluorescence monitoring revealed that the cells retained viability and exhibited more efficient expression of P2X3 at 30 °C than at 37 °C (Figure S1C, D). A calcium influx assay indicated that P2X3-GnTI^-^stable cell lines expressed functional P2X3 (Figure 1A).

**Figure 1.**
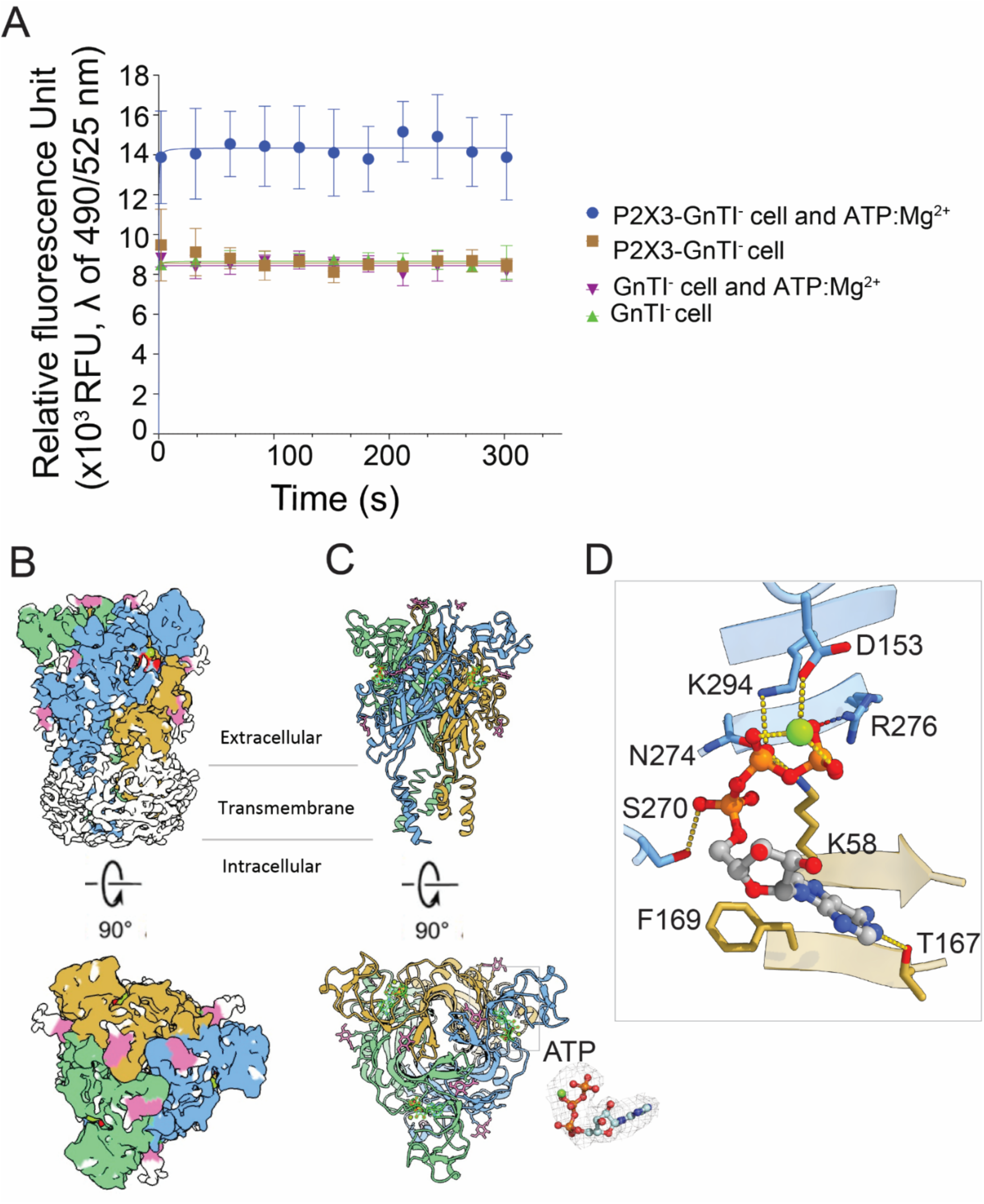
The Expi293F GnTI-stable cell line expressing functional P2X3 receptor. (A) Intracellular Ca^2+^ signaling in GnTI^-^ and P2X3-GnTI^-^ stable cell lines are compared. The cells were loaded with the Ca^2+^-specific indicator Fluo-8 and stimulated with ATP:Mg^2+^. Error bars represent standard deviation, with n=3-6 for each test conducted. (B) The overall structure of the P2X3:ATP complex reconstituted in DDM. Each subunit color-coded is presented with the protomers of the trimer-colored light blue, light orange, and green. Representations of the cryo-EM map viewed parallel to the membrane (top panel) and perpendicular to the membrane from the extracellular side (bottom panel). (C) The structure is depicted as cartoons. ATP bound to the receptor is represented as stick-spheres, viewed parallel to the membrane (top panel) and perpendicular to the membrane from the extracellular side (bottom panel). The electron density map for ATP contoured at 3.0 σ. (D) A close-up view of the ATP:Mg^2+^ binding site is provided in the inset. ATP bound to the receptor is represented as stick-spheres. Mg^2+^ is represented as a green sphere.

### Binding of Camlipixant and ATP to P2X3 receptor

To establish a complex between P2X3 and camlipixant, we first purified the receptor (Figure S2A) and assessed its binding affinity to camlipixant using Isothermal Titration Calorimetry (ITC). However, no binding of camlipixant to P2X3 was observed. We hypothesized that extracellular ATP released during cell lysis binds to the receptors, as previously observed in insect cells, and blocks the camlipixant-binding site (Mansoor *et al*, 2016). Our cryo-EM structure confirmed the presence of ATP:Mg^2+^ bound to P2X3 (Figure 1B-D, Figure S3, Table S1), which closely resembled the previous X-ray crystal structure of P2X3. Superposition of the refined models obtained from crystallography and cryo-EM showed that the Cα atoms overlapped with a Root Mean Square Deviation (RMSD) of 0.81 Å (PDB-ID 5SVK) (Figure S4). To remove the bound ATP, we carried out extensive dialysis of the P2X3:ATP complex in a buffer containing 20 mM Tris-HCl (pH 7.6), 1000 mM NaCl, and 0.08% DDM, supplemented with a protease inhibitor, for 7 days. Purification of the dialyzed protein, followed by binding studies, showed that P2X3 bound to camlipixant with a *K_d_* of 100 nM using ITC (Figure 2A, Figure S5). These data indicate that camlipixant binds to pockets exposed to the ATP-unbound state (also called the resting state). Intriguingly, we also observed that the camlipixant-bound receptor was incapable of binding to ATP:Mg^2+^, suggesting that camlipixant occluded the ATP-binding site (Figure 2B, Figure S6). The binding affinity of ATP to P2X3, measured by MicroScale Thermophoresis (MST) is 2μM (Figure 2B, Figure S6). There are reports in the literature of this binding affinity varying from nanomolar to micromolar (Vitiello *et al*, 2012).

**Figure 2.**
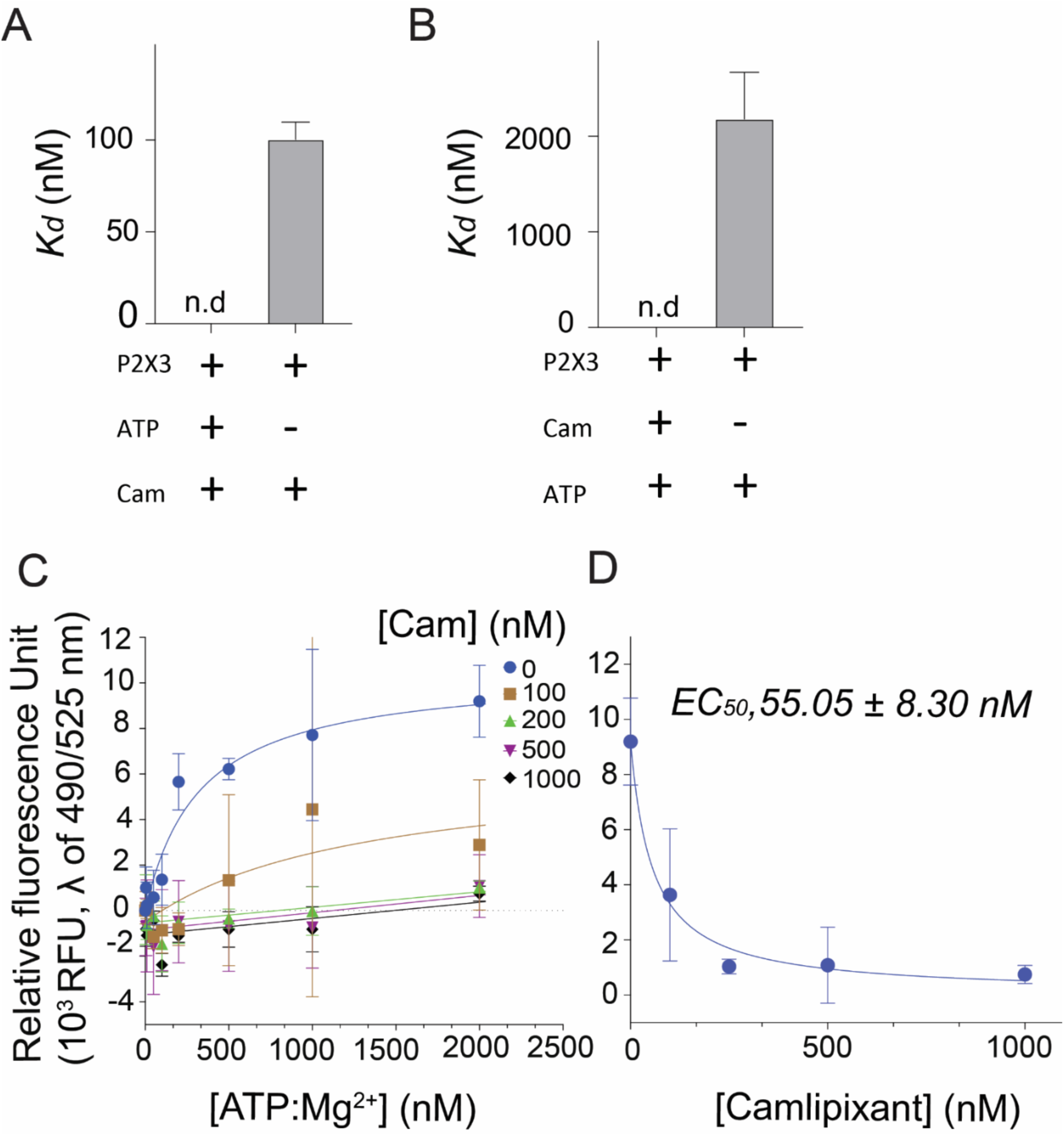
Inhibition of P2X3 activity by camlipixant. The binding of either ATP or camlipixant (Cam) to the P2X3 receptor prevents another binding event. (A) When P2X3 was incubated with ATP there is no binding of camlipixant; in the absence of ATP, camlipixant binds with K_d_ of 100 nM. (B) When P2X3 was incubated with camlipixant ATP did not bind, however in camlipixant’s absence ATP binds with a K_d_ of 2 μM. (C). Camlipixant’s inhibitory effects on intracellular calcium flux by ATP:Mg^2+^ on the P2X3-GnTI^-^ cell line were measured. The change in intracellular Ca^2+^ was monitored for 5 minutes, with baseline measurements taken 1 minute before ATP:Mg^2+^ addition. There is a reduction in calcium uptake with increasing concentrations of camlipixant. (D) P2X3-GnTI- cells were incubated with different concentrations of camlipixant for 30 minutes. Fluorescence was measured 5 min after addition of 2 μM ATP:Mg^2+^. Error bars represent standard deviation, with n=3 for each test conducted. n.d, not determined.

To elucidate the molecular mechanism of inhibition by camlipixant, we assessed the dose-response of P2X3-mediated intracellular Ca^2+^ influx in the presence or absence of various concentrations of camlipixant and ATP:Mg^2+^ in P2X3-GnTI cells. Concentrations exceeding 2 μM ATP:Mg^2+^ induced fluorescence in P2X3-GnTI^-^ cells in a dose-dependent manner (Figure 2C). Notably, camlipixant-treated P2X3 receptors exhibited a dose-dependent decrease in fluorescence upon addition of ATP:Mg^2+^ (Figure 2C). Camlipixant inhibited ATP-activated P2X3-mediated Ca^2+^ ion transfer in a concentration-dependent manner, with an IC_50_ of 55.05 nM (Figure 2D).

### Overall structure of camlipixant-bound P2X3 reconstituted into peptidisc

To elucidate the structural basis of camlipixant binding, our initial endeavor was to determine the cryo-EM structures of the P2X3 receptor bound to camlipixant. The experimental conditions mirrored those for P2X3:ATP in the presence of DDM detergent. However, we encountered challenges related to preferred orientation. Although we obtained a high-resolution map of P2X3:camlipixant (2.94 Å resolution), we were unable to resolve the transmembrane domains (Figure 3A, Figure S7A-D).

**Figure 3.**
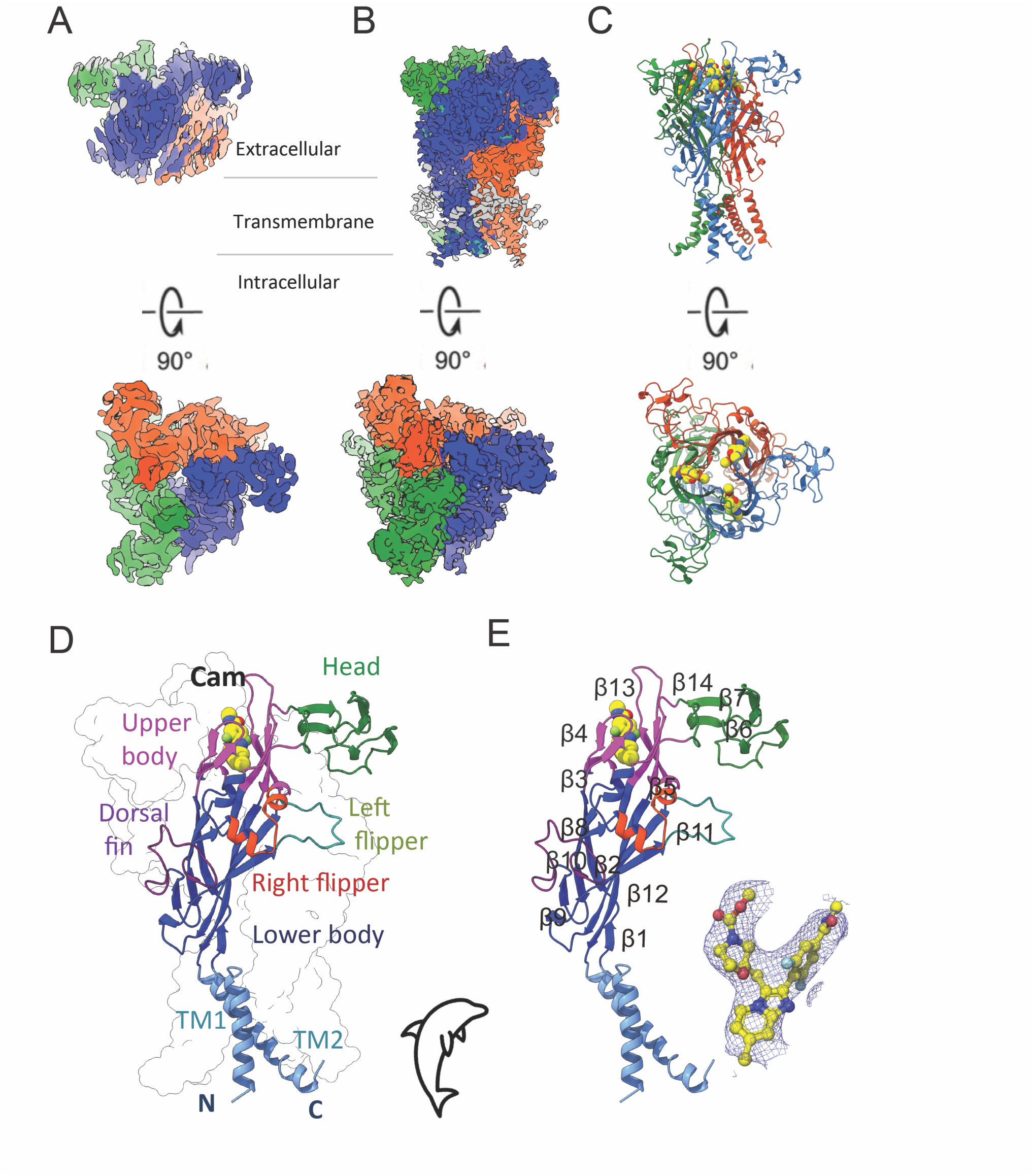
Cryo-EM structures of the camlipixant-bound P2X3 receptor, reconstituted into either DDM or peptidisc. Representation of the cryo-EM map of the Cam-bound P2X3 receptor reconstituted into DDM (A) or peptidisc (B), shown in side view and top view. (C) The overall structure of the P2X3:camlipixant complex reconstituted into peptidisc, with each subunit color-differently, is depicted. The protomers of the trimer are colored blue, orange, and forest. (D, E) The P2X3 monomer adopts a dolphin-like shape, with each region of the P2X3 protein depicted in colored cartoon representation according to this shape. Camlipixant (shown as spheres) binds to the upper body domain of P2X3. The electron density map for camlipixant is also shown and contoured at 3.0 σ. The beta-sheet numbering is continuous from the N-terminus to the C-terminus.

To resolve the full-length receptor complexed with camlipixant, we employed the peptidisc approach to stabilize the receptor and bind it to camlipixant (Angiulli *et al*, 2020; Couston *et al*, 2023; Carlson *et al*, 2018). Notably, reconstitution into the peptidisc significantly improved the distribution of the receptor onto the cryo-EM grid compared with reconstitution in the DDM detergent (Figure S7E, F). A high-quality cryo-EM dataset of P2X3:camlipixant reconstituted into a peptidisc was collected and processed at a resolution of 3.6 Å (Figure S8, Table S1). The map was further refined by density modification, resulting in an overall resolution of 3.4 Å (Figure 3B, S8). The cryo-EM density exhibited particularly well-defined features, with a local resolution ranging from 3 to 3.2 Å resolution (Figure S8). We unambiguously rebuilt and assigned side chains to almost all residues of the transmembrane helices (Figure 3C and S9). We modeled residues 21–345 from our initial construct, indicating that only the first 15 and last 18 amino acids in the C-terminus (which included a linker) were not observed. The extracellular domain was well defined, as evidenced by the local resolution map (Figure S8). Another indication of cryo-EM map quality was the presence of additional density surrounding the P2X3 transmembrane helices, suggesting the presence of peptidiscs and small lipids in the detergent. The peptidisc was arranged in two to three layers, perpendicular to the helices of the transmembrane domains (Figure 3B). However, these helices were not built into the final model deposited in the Protein Data Bank (PDB), because the side chains and helix orientations could not be confidently assigned, as previously described (Carlson *et al*, 2018b; Angiulli *et al*, 2020).

P2X3 reconstituted in the peptidisc formed a trimeric receptor, which is consistent with all previously reported P2X3 structures (Figure 3B, C). The structure can be divided into three main domains: extracellular, transmembrane, and intracellular. To validate the peptidisc strategy as a robust alternative for investigating P2X3 structure, it is imperative to assess the preservation of its structural integrity under these conditions. A structural comparison with the X-ray crystallography-determined structure revealed that the P2X3 structure inserted into the peptidisc was comparable. Superimposing our structure onto P2X3 (PDB-ID, 5SVJ) in DDM, suggested high structural similarity, with RMSDs of 0.92 Å overall and 0.50 Å in transmembrane helices (Figure S10A, B). This observation strongly suggests that peptidisc stabilization preserves the structural integrity of P2X3 because no major structural rearrangements were observed.

### A cryptic drug binding site in P2X3

In short, a single subunit of the P2X3 receptor displays a distinctive ‘dolphin-like’ shape, akin to the broader P2X receptor family (Figure 3D, E). Notably, camlipixant binds within the pocket formed by the upper body domain between adjacent subunits positioned above the orthosteric binding site, that is, the ATP-binding pocket (Figure 3D, E). This drug-binding pocket is encircled by 20 residues, predominantly protruding from the β-strands (β3, β4, β5, β13, and β14) within the upper body domains of neighboring subunits (Figure 3E, Figure 4A, B, Table S2). At the current local resolution of approximately 3 Å, the distances and angles between the side chains and drug molecules can be discerned (Figure 4A, B, Figure S8).

**Figure 4.**
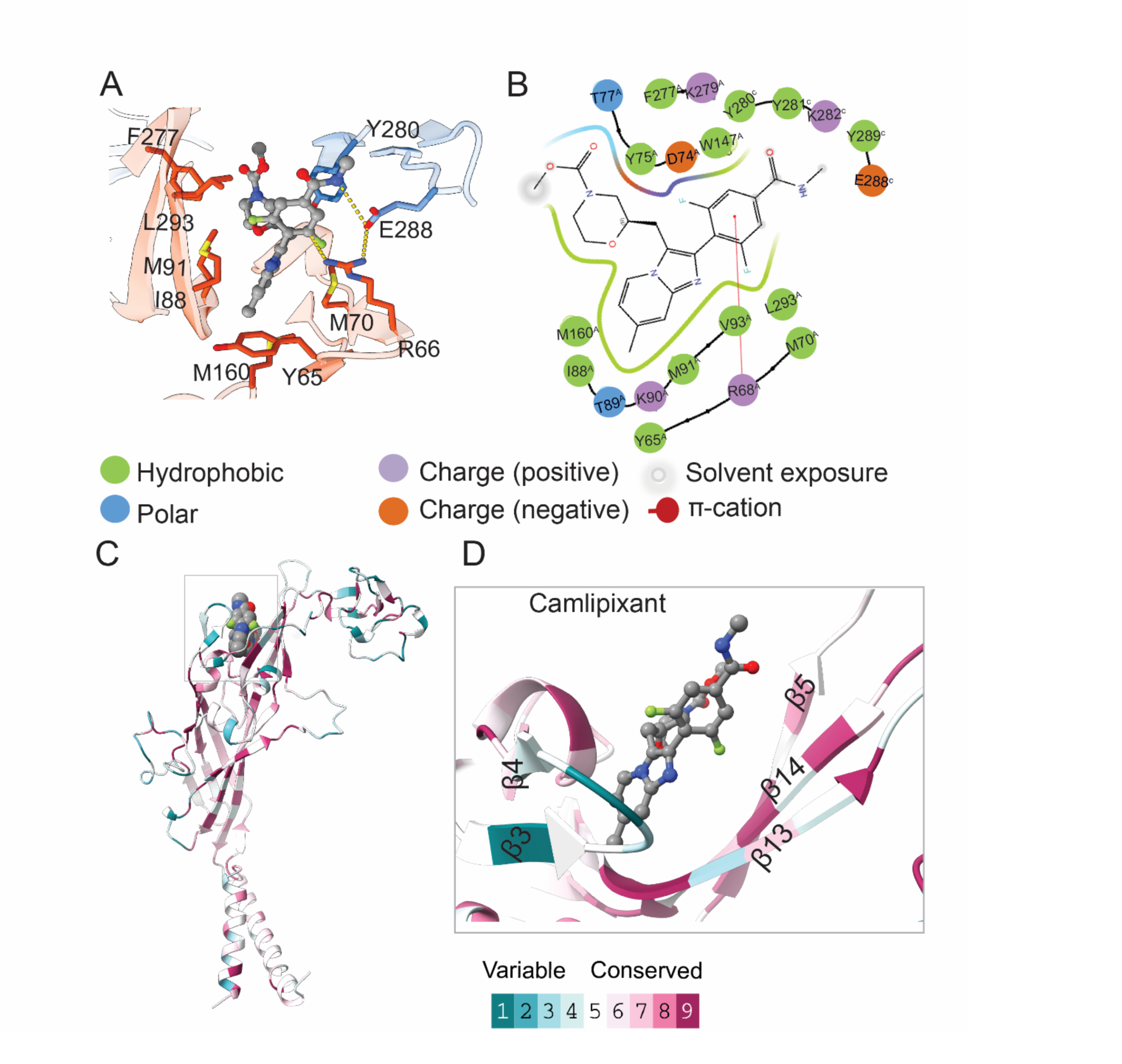
The binding pocket for camlipixant in the P2X3 receptor exhibits moderate conservation among P2X receptors. (A) A close-up view of the camlipixant binding site in P2X3. (B) A schematic diagram illustrating the interactions between Cam and P2X3. (C) Residues involved in camlipixant binding show moderate conservation across P2X receptors. The evolutionary conservation of residue positions on the P2X3 structure is estimated based on phylogenetic connections among homologous sequences. (D) A close-up view of the camlipixant binding site highlighting conserved residues.

The extensive hydrophobic interactions observed between the receptor and camlipixant underscored significant aspects of their binding dynamics. Specifically, the involvement of residues M70, M91, F277, and L293 in these interactions highlights their pivotal role in stabilizing the receptor-camlipixant complex. The involvement of multiple residues, such as phenylalanine (F) and leucine (L), indicates a complex network of intermolecular contacts contributing to a strong preference for nonpolar environments, suggesting that the binding interface between the receptor and camlipixant is largely hydrophobic. Additionally, R68 and E288 recognize camlipixant by cation-ρε and charge-charge interactions with N04 and the difluorophenyl ring, respectively, acting as a lid in the binding process (Figure 4A, B). The evolutionary conservation of positions of residues in the P2X3 structure was estimated based on the phylogenetic connections among homologous sequences using Consurf (Glaser *et al*, 2003), suggesting that these camlipixant-bound residues are moderately conserved in P2X3-7, but distinct from those found in P2X2 receptors. (Figure 4C, D, S11, and S12 and Table S2).

### Camlipixant binding narrows down ATP binding pocket size

Upon camlipixant binding to the receptor, conformational changes occur in P2X3. The structure of P2X3 in the absence of antagonists (apo-form) showed a closed conformation, as evidenced by constriction of the channel gate by transmembrane helices at residues I318 and V321 (Figure 5A). The structural similarity of the transmembrane helices between the camlipixant-bound P2X3 structures and the apo-state represents a closed conformation (PDB-ID, 5SVJ), indicating that camlipixant likely stabilizes the resting closed state (Figure 5A-C, Figure S10A, B, Video1). During camlipixant binding, there is a minor change in the “upper body” domain, where the residues mainly projecting from β-strands (β3, β4, β13, and β14) move backward by approximately ∼3 Å, enlarging the camlipixant binding pocket with a volume increase from 513 to 1203 Å^3^ (Figure S10A, B, Video1). Additionally, the ATP-binding pocket loop moves closer, narrowing the pocket volume from 1161 Å^3^ in apo to 573 Å^3^ in the camlipixant-bound state (Figure S10A, B, S13B-C, Video1). This alteration prevents ATP-mediated receptor activation of P2X3 upon binding to camlipixant. Significant conformational changes are observed in P2X3 between the ATP-bound and camlipixant-bound states. The “upper body” domain moves backward by approximately 10 Å, enlarging the pocket for camlipixant binding approximately fivefold (241 versus 1203 Å^3^), while the ATP binding pocket is narrowed down around two-fold in the camlipixant-binding state (1143 vs 573 Å^3^) (Figure S10 C, D, S13A-C, Video 2). Additionally, the transmembrane helices moved closer to the gate (Figure 5A, B, D, Figure S10C-D, Video 2). Consistent with the activation mechanisms proposed for P2X3 and P2X4, ATP binding brings the dorsal fin domain towards the head domain and pushes the left flipper domain away from the ATP-binding pocket (Mansoor *et al*, 2016; Kasuya *et al*, 2017; Shen *et al*, 2023). These movements are coupled with the widening of the lower body domain, albeit to a degree where the transmembrane helices remain closed (Figure S10, Video 2). Interestingly, we also observed changes in the turret volume during the Cam-binding state, which is closed in the ATP-bound state, partially open in the apo state, and fully open in the camlipixant-bound state (Figure 5, Figure S13D-F). Therefore, our data suggest that the drug-binding pocket narrows upon ATP binding and that such conformational rearrangement is crucial for P2X3 channel opening. Conversely, ATP release induces conformational changes that open the drug binding pocket. Camlipixant binding narrows the ATP-binding pocket, which is essential for inactivating P2X3-dependent ATP signaling (Figure 5 and S10).

**Figure 5.**
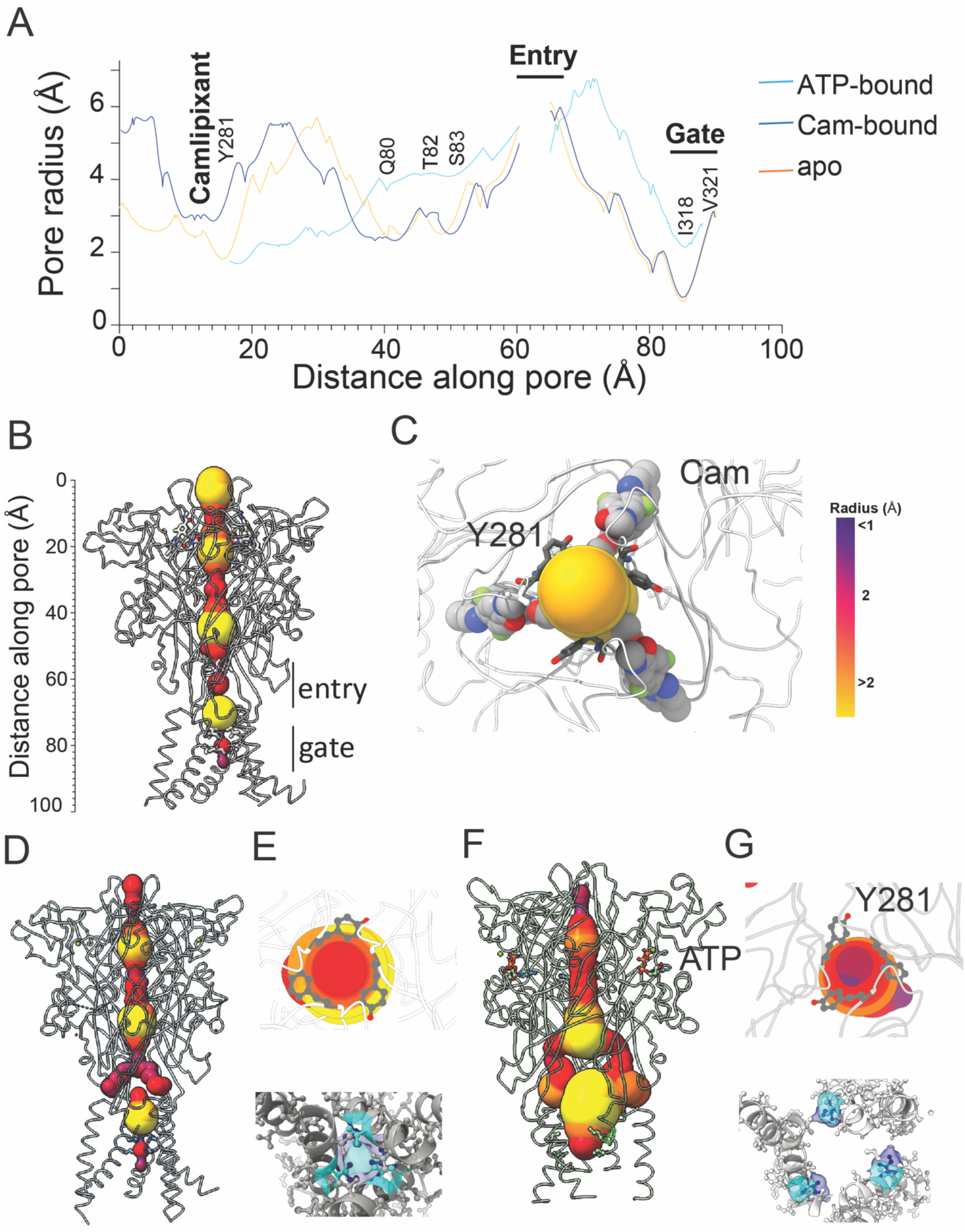
The turret of the P2X3 receptor opened, but the ion entry and gate remained closed in camlipixant-bound state. (A) The channels formed in camlipixant-bound, apo- (PDB-ID, 5SVJ), and ATP-bound states of P2X3 are represented by line diagrams. Residues lining the selectivity filter and surrounding the gate are highlighted. (B, C) Surface representations of the channels from camlipixant-bound P2X3 are viewed from two perpendicular orientations with the respective camlipixant and Y281 residue involving in turret formation. (D-G) Surface representations of the channels from Apo (D, E) and ATP-bound state (F, G) with the respective Y281 residue involving in turret formation. The gate is formed by residues I318 (blue) and V321 (cyan) viewed from intracellular to extracellular (bottom view).

### The ion entry and gate are closed during camlipixant-bound state

To investigate whether allosteric inhibition by camlipixant prevents narrowing of the turret upon ATP binding in P2X3, we utilized whole-channel calculations. The entire channel, approximately 85 Å in length, was divided into two parts: the turret and the transmembrane channel separated by ion entry (Figure 5A). The ion entry and gate were widened in the ATP-bound state but closed in both the apo- and camlipixant-bound states (Figure 5A). Notably, the channel analysis in the apo- or camlipixant-bound state did not reveal any significant changes in the entire channel (Figure 5A, B, S10A, B, video 1). Conformational changes in the upper-body domain, including the turret and Cam-binding pocket, support the notion that turret opening is essential for P2X3 channel closure, whereas turret closure leads to channel opening (Figure 5B, C; Figure S10, S13, video 1,2).

Adjacent to the upper ion entry, the turret is closed in both the ATP-bound and apo states but opened in the camlipixant-bound state, resulting from conformational changes in residues Q80, T82, and S83 (Figure 5A, D, F). Camlipixant binding disrupts conformational changes in the upper body domain, causing insufficient movement of the pore-lining transmembrane helices to simultaneously open the channel. Overall, our data suggest that the P2X3 receptor undergoes conformational rearrangements, where both the camlipixant-binding pocket and the turret in the P2X3 receptor widen upon drug binding (Figure 5B, C, Figure S13). As such conformational changes are required to prevent ATP binding and stabilize channel closure, camlipixant binding to P2X3 prevents these constrictions, thereby efficiently blocking receptor activation (Figure 5 and S13). Furthermore, in both the apo and camlipixant-bound states, the gate’s radius diminishes from 2 Å to 0.7 Å, predominantly influenced by crucial residues, I318 and V321(Figure 5A, E, G). This alteration signifies the obstruction of cation transfer, encompassing Ca^2+,^ Na^+^, and K^+^ with ionic radii of 0.99, 0.95, and 1.33 Å.

### The camlipixant binding site appears highly selective in P2X3 receptor

The crucial residues within the camlipixant-binding pocket differ between the P2X3 and P2X2 isoforms, as depicted in Figure S12 and Table S2. This reinforces the highly selective targeting of P2X3 over P2X2/3 receptors by camlipixant (Field, 2023; Garceau & Chauret, 2019). Utilizing our recently resolved camlipixant-bound P2X3 complex structure, we modeled both the P2X2 structure and the P2X2/3 heterotrimeric form binding to camlipixant (Figure 6). Given the plethora of human P2X2 receptor isoforms, we chose isoform K because of its high sequence identity with P2X3 (Figure S12B). Notably, we observed discrepancies not only in the sequence of the camlipixant-binding site, but also in the structure of the camlipixant-binding pocket itself. Specifically, the β13 and β14 regions shifted forward towards the adjacent subunit, thereby narrowing the binding pocket by approximately 5%, potentially inducing significant alterations in the camlipixant binding pocket (Cam1; Figure 6A, B). Our analysis also revealed that the loop between β4 and β5 in P2X2 is longer and clashed with β13 and β14 in the adjacent subunit when bound to camlipixant (Cam2; Figure 6A, C). Furthermore, we used the P2X2/3 structure from the camlipixant-bound state as the initial conformation and started MD simulations. Trajectories were analyzed to assess ligand stability and protein conformational changes. The lowest binding free energy was obtained after a 100 ns conformational sampling process that explored all potential binding modes of the receptor and ligand conformations (Figure S14, Table S3). We noted a considerable decrease in the MMGBSA binding free energy at both binding interfaces between P2X2-P2X3 (Cam1) and (Cam2) subunits compared to the camlipixant-bound P2X3-P2X3, −68.25 ± 4.64 and −47.58 ± 4.69 kcal/mol respectively, versus to - 74.53 ± 3.32 kcal/mol (Table S3). The camlipixant is mostly bound to the outside of the upper vestibule; the hydrophobic group is exposed to a solvent environment during the simulation, indicating that camlipixant could not bind well to the P2X2/3 receptor interface (Figure 6B, C).

**Figure 6.**
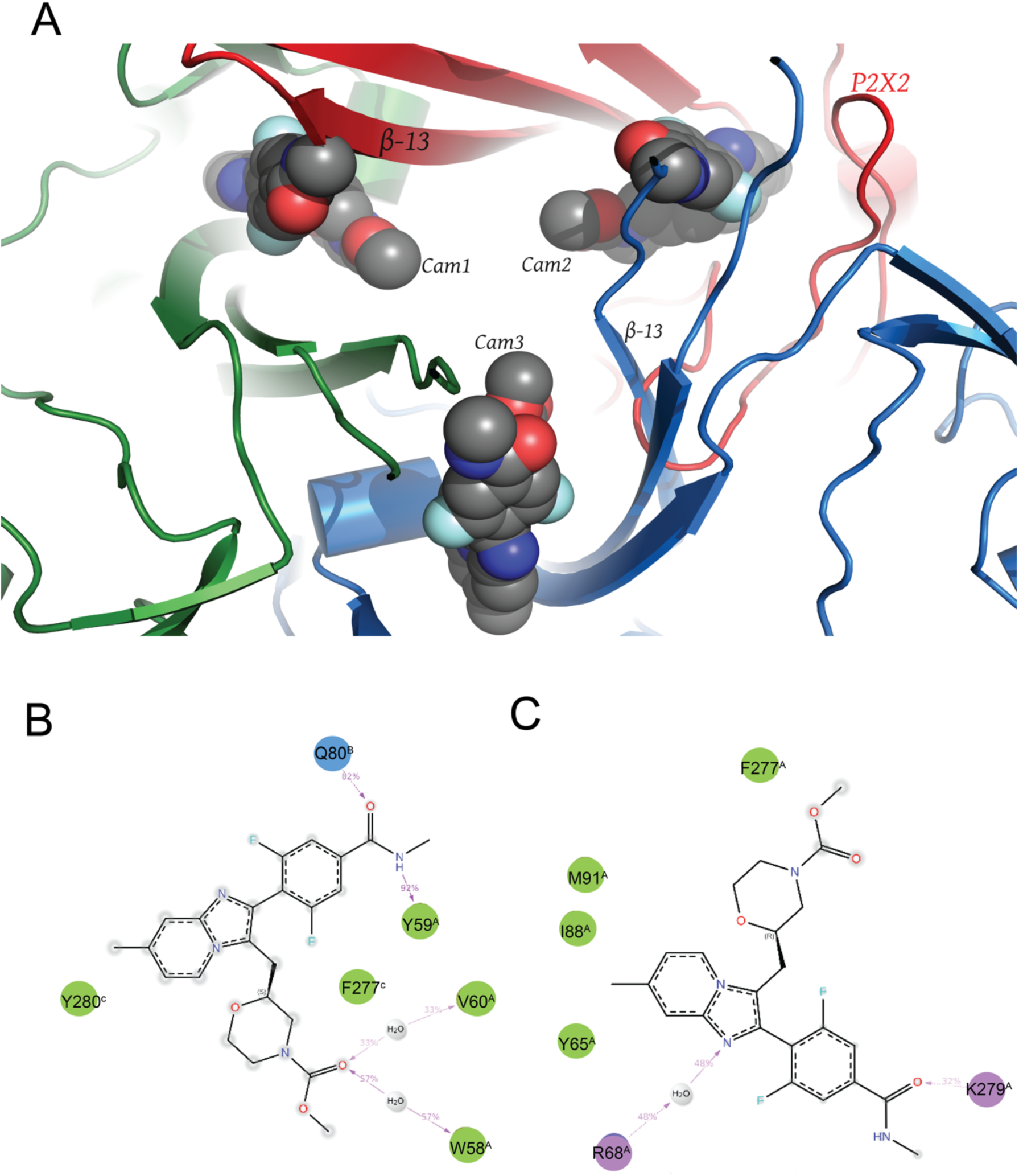
Camlipixant exhibits selective targeting of the P2X3 receptor over the P2X2/3 receptor. (A) Camlipixant demonstrates preferential targeting of the P2X3 receptor over the P2X2/3 receptor. (A) The illustration depicts a model of the P2X2/3-camlipixant (Cam) complex, wherein the P2X2/3 receptor is overlaid onto the P2X3 receptor shown in a cartoon diagram. The camlipixant ligand is represented as spheres. Cam1 and Cam2 are situated at the interface of P2X2 (red, chain A), while Cam3 binds within the interface of P2X3. (B, C) A schematic diagram outlines the interactions between Cam1 and Cam2 with P2X2/3 post molecular dynamics simulations of the receptor bound to the heterotrimer.

*This binding site demonstrates a notable preference for P2X3 over P2X2 isoforms, corroborating earlier experimental findings that highlight the precise targeting of P2X3 while avoiding taste loss linked with P2X2/3 receptors (Garceau & Chauret, 2019)*. Conversely, gefapixant (formerly known as AF-219 or MK-7264), which targets a distinct conserved allosteric binding site, has shown considerable therapeutic efficacy in clinical trials for the treatment of refractory chronic cough (Abdulqawi *et al*, 2015; Wang *et al*, 2018). The binding site is encompassed by regions such as the dorsal fin, left flipper, and lower body domains of the two adjacent subunits, forming a pocket in close proximity to the orthosteric site (Figure S15). Examination of the gefapixant binding pocket revealed highly conserved sequences and structures between P2X2 and P2X3 (Figure S12 and S15) with RMSD values of 0.3 Å. This binding site exhibits strong connectivity between P2X3 and P2X2/3 receptors (Figure S15), consistent with data suggesting that gefapixant is linked to taste loss as a side effect (Field, 2023; McGarvey *et al*, 2022). Collectively, our structural analyses affirm that the newly identified cryptic binding pocket exhibits high selectivity for P2X3 over P2X2 compared to the original allosteric binding pocket on the P2X3 homotrimer receptor.

Additionally, the inter-subunit cavity formed by β13 and β14 in the upper body domain is notably wider in P2X3 than in Zebrafish P2X4 and Panda P2X7, as reported previously (Jiang *et al*, 2021; Shen *et al*, 2023). The measured volume of the binding pocket is 1203 Å^3^ for P2X3, 1034 Å^3^ for P2X4, and 1183 Å^3^ for P2X7. The equivalent pocket in the P2X4 and P2X7 receptors is narrow enough to accommodate camlipixant, despite its hydrophobicity being similar to that of the P2X3 receptor. Although the structures of P2X5 and P2X6 have not been reported, the volume of the binding pocket differs from that of P2X3 because of variations in the key residues (Figure S12, Table S2). The ‘turret-like’ structure and the cleft corresponding to the P2X3 camlipixant-binding pocket remain relatively occluded in the P2X3 receptor (Figure S16). These findings support the notion that the narrowing of the inter-subunit space appears to be unique to the P2X3 receptor. Therefore, differences in the sizes of the inter-subunit hydrophobic pocket and antagonist drugs are crucial for conferring P2X3-selective binding of inhibitors.

To investigate the potential cross-targeting of camlipixant and other antagonists of P2X3, P2X4, and P2X7, we performed flexible docking interactions with P2X receptors, and identified these interactions during MD simulation. Our data indicate that camlipixant exhibits binding affinity to P2X3 over P2X4 and P2X7, as evidenced by the lower free energy values and docking score of −8.35, −7.31 and −6.00, respectively (Figure 3E, Figure S16). The values of MMGBSA binding free energy between camlipixant to P2X7, P2X4 are −61.92 ± 3.72 and −72.54 ± 4.32 kcal/mol respectively, versus to −74.53 ± 3.32 kcal/mol inn that of P2X3 (Table S3). The hydrophobic group is inserted into the hydrophobic lumen inside the pocket of the P2X3 receptor, and the carboxyl group mainly interacts with the charged residues, whereas there is less interaction between the hydrophobic and hydrogen bonds between the P2X4 receptor and camlipixant. The carboxyl group of camlipixant is mainly exposed to the hydrophilic solvent outside the pocket, while there is potential solvent exposure in the interaction between the P2X7 receptor and camlipixant (Figure S16C-E, Figure S17). Collectively, our data suggest that camlipixant potentially binds to P2X4 and P2X7 in similar binding pockets, albeit with lower affinity.

## Discussion

The specific localization of P2X3 receptors in primary sensory neurons plays a crucial role in modulating synaptic transmission, inflammation, and pain sensation, including neuropathic pain, migraine, and visceral pain disorders, rendering P2X3 an attractive target for therapeutic interventions. P2X3 receptors are crucial for sensory neuron function in various tissues and organs. Their significance is particularly pronounced not only in the cough reflex but also in pain perception and bladder function (Ford & Undem, 2013). The development of drugs that target P2X3 receptors is a crucial pursuit in clinical research. Nonetheless, achieving subtype selectivity is imperative to minimize the toxic side effects and enhance the potential for successful drug development. Despite the significant pharmaceutical interest in targeting P2X3, no drug has received FDA approval to date. Therefore, chronic cough, chronic obstructive pulmonary disorder, and bladder disorders remain unmet clinical needs.

There has been extensive interest in the search for molecules that act as selective antagonists of P2X3 over the P2X23 and P2X receptors. Among the numerous new antagonists targeting P2X3 that have emerged in recent years, camlipixant is currently undergoing Phase III clinical trials (Garceau & Chauret, 2019; Davenport *et al*, 2021; Field, 2023). The efficacy and selectivity of the camlipixant were evaluated across various P2X receptors using cell-based calcium mobilization assays following activation by ATP (Garceau & Chauret, 2019). Camlipixant exhibited potent and non-competitive antagonistic effects on hP2X3 receptors, both in the cloned form within mammalian cell lines and in native P2X3 receptors from primary neurons of the human dorsal root ganglia, with an IC_50_ of approximately 20 nM. Similar potency was observed against the P2X3 receptor variant. Conversely, camlipixant did not demonstrate significant antagonistic activity against hP2X1, hP2X2b, hP2X4, hP2X5, and hP2X7 receptors, with EC_50_ values in the µM range (Garceau & Chauret, 2019).

The cryo-EM structure of a P2X3-camlipixant complex identified a cryptic drug-binding site for camlipixant in the P2X3 receptor. Through structural analyses and biochemical and biophysical experiments, we found that camlipixant binds to accessible pockets present in the resting state, rather than solely to sites exposed when the channel is open, such as those deeper within the upper body domain. Camlipixant-binding induces conformational changes in the extracellular domain machinery that block the ATP-binding site and stabilize the closed ion entry gate (Figure 7). Interestingly, the size of the closed gate completely blocked ion transfer, suggesting potential implications for neuronal signaling (Figure 5). The druggable site is situated within the interface created by β-strands 3, 4, and 13, 14 in the upper body region of the P2X receptors. Sequence identity is moderately conserved across the P2X family (Table S2). Additionally, the identified pocket is distinct from that targeted by the FDA-rejected drug gefapixant/AF-219, which is formed by the left flipper and dorsal fin (Figure 3, S15). These binding sites provide significant selectivity to the P2X3 homotrimer rather than the P2X2/3 heterotrimer, which provides a molecular explanation for the physiological observation of not dampening the taste response while still reducing the cough response.

**Figure 7.**
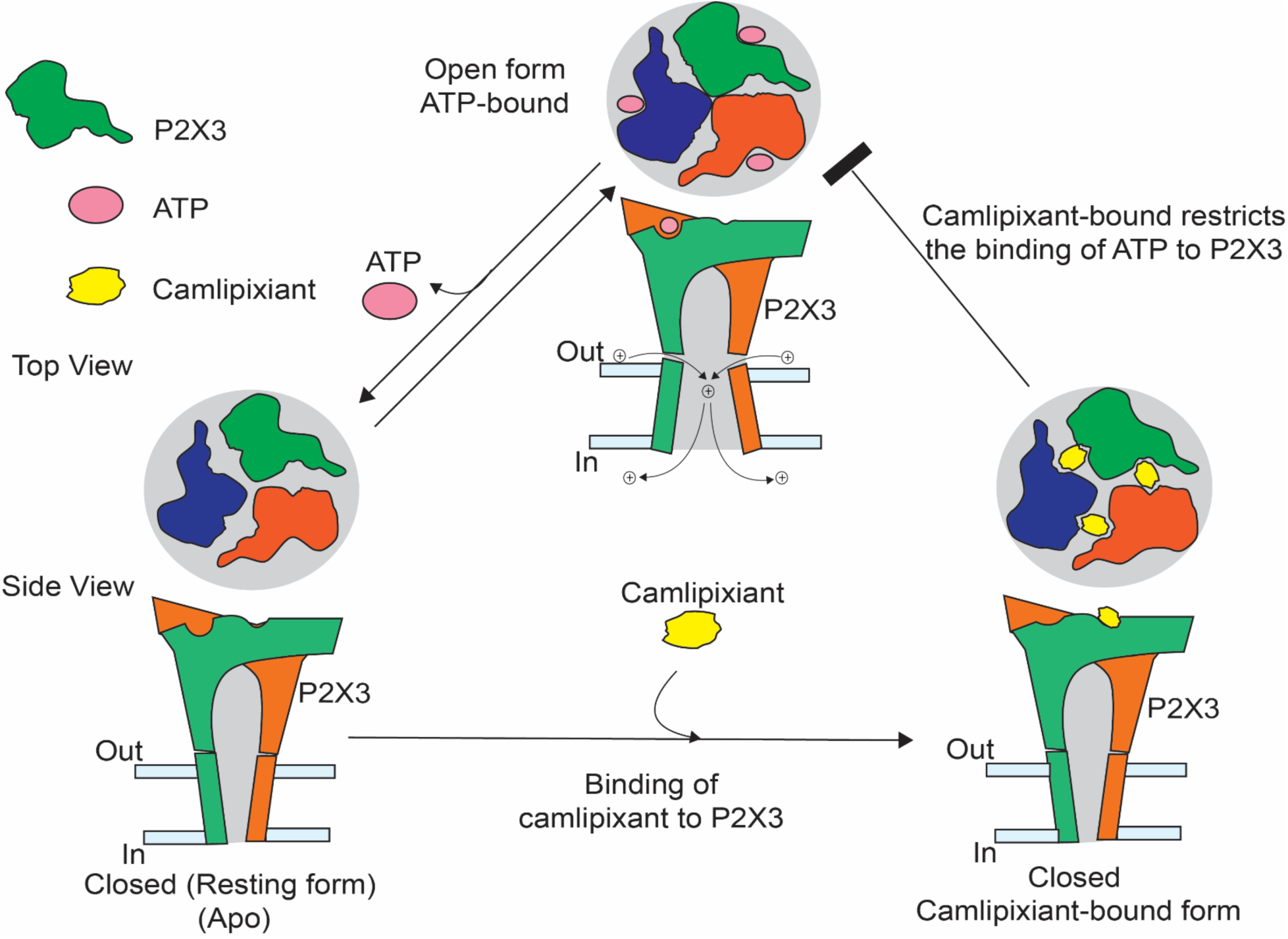
Mechanisms underlying camlipixant-bound P2X3 inhibition. Schematic representations of the P2X3 receptor, viewed from the top and side, display distinct subunits in various colors. As the channel activates, the drug-binding pocket narrows, and the ion entry gate opens. Upon ATP dissociation, a conformational change occurs in the receptor, enlarging the drug-binding pocket to facilitate drug binding; ion entry and gate opens. Camlipixant, selective to P2X3, stabilizes the closed conformation by obstructing the movement of inter-subunit cavities, ATP binding site closes.

## Materials and Methods

### P2X3 construction, expression and purification

The gene coding for P2X3 from *Cannis lupus* sp. (cP2X3) was cloned into the pElanco3 vector with the deletion of 5 first residues in the N-terminal and the 33 last residues in the C-terminal, as previously described in human P2X3 (Mansoor *et al*, 2016). The gene construct expressed the 12His-eGFP-GGS linker-Thrombin Cleavage site-GGS linker-Protein-GGS-HRV3c Cleavage site-Rho1E tag with codon optimization for the HEK cell line. To generate the P2X3-Expi293F GnTI^-^ suspension cell line, the pElanco3-eGFP-P2X3 Transposon and PiggyBac Transposase plasmids were transfected at a 1:20 ratio of transposase:transposon using a Neon Electroporation System. Briefly, 5 x 10^7 cells were transfected with 10 µg of total DNA using the Neon 100 uL tip 6-well plate protocol. Neon tips (100 µL) were filled with the cell and plasmid mixture, and the cells were electroporated with two pulses of 1,100 V each, with a pulse width of 20 ms, and then transferred into 2 mL of pre-warmed Gibco Expi293 Expression Media (Cat# A1435101) in a low-attachment 6-well plate and incubated at 37 °C in a humidified 8% CO_2_ incubator for one day. The expression of the receptors in these vectors is driven by constitutive promoters. After 24 h, GFP expression was verified by fluorescence microscopy and 2 µg/mL puromycin was added to initiate cell selection. After 7 days of puromycin selection, the cells were expanded in Corning Erlenmeyer shake flasks in Gibco Expi293 Expression Media supplemented with 1 µg/mL puromycin, and then frozen in Expi293 Expression Media supplemented with 10% DMSO.

P2X3 was isolated and purified from the stable cell line P2X3-Expi293 GnTI^-^ using Expi293™ Expression Medium supplemented with 2 µg/ml puromycin (Thermo Fisher, Cat#, A1113803). The cells were cultured at 37 °C in an 8% CO_2_ atmosphere and shaken at 125 rpm. Once the cell density reached approximately 2 x 10^^6^ cells/ml, the temperature was lowered to 30 °C and maintained for six days. The cells were harvested, washed with phosphate-buffered saline (PBS; pH 7.4; Gibco) via centrifugation at 6,000 rpm, flash-frozen, and stored at −80 °C until further use.

The cells were disrupted using a sonicator, operating at 45% amplification with a pulse of 10s on and 10s off, in the presence of buffer A (20 mM Tris-HCl pH 7.6, 150 mM NaCl, 0.1 mM TCEP, 5% glycerol) supplemented with an EDTA-free protease inhibitor cocktail (Thermo Fisher, Cat # PIA32955). Undisrupted cells were removed by centrifugation at 4,000 ×rpm for 10 min at 4 °C. The resulting suspension containing disrupted cells and membranes was harvested and subjected to ultracentrifugation at 40,000 ×rpm for 1 h at 4 °C. The membrane-containing pellet was homogenized using a homogenizer in buffer B (20 mM Tris-HCl pH 7.6, 150 mM NaCl, 0.1 mM TCEP, 2% DDM) supplemented with a protease inhibitor cocktail. After rotating the homogenized solution for 2 h at 4 °C, it was subjected to ultracentrifugation. The soluble fraction obtained was applied to Talon beads ( Cat # 635503; Takara) and incubated for 6 h at 4 °C. Subsequently, the beads were washed with buffer C (20 mM MES pH 6.5, 150 mM NaCl, 0.1 mM TCEP, 0.2% DDM) before treatment with Thrombin and HRV3C protease in the same buffer for 6 h at 4 °C. The P2X3 fraction was then harvested and subjected to size-exclusion chromatography (Superdex 200 10/300 column) in the presence of buffer D (20 mM Hepes pH 7.5, 100 mM NaCl, 0.08% DDM). Peak fractions containing P2X3 were pooled and analyzed by SDS-PAGE. Monodisperse fractions of P2X3 were used for cryo-EM grid preparation.

### Camlipixant-bound P2X3:peptidisc reconstitution

To dissociate ATP from the P2X3 receptor, the sample was subjected to dialysis in a buffer containing 20 mM Tris-HCl (pH 7.6), 1000 mM NaCl, and 0.08% DDM supplemented with protease inhibitor for 7 days. The buffer was refreshed daily before returning it to buffer D.

The peptide sequence was chemically synthesized using solid-phase peptide synthesis (GenScript) to achieve a purity of > 85%. The peptide was dissolved in buffer E (20 mM HEPES pH 7.5, and 100 mM NaCl) to a final concentration of 1 mM. To reconstitute P2X3 into the peptidisc, a sample of purified P2X3 in DDM was incubated for 30 min with an excess of peptidisc solution at a 1:20 molar ratio in the presence of buffer E (20 mM HEPES pH 7.5, 100 mM NaCl) on ice. The P2X3:peptidisc complex was separated from the free peptidisc by size-exclusion chromatography (SEC) using a Superdex 200 Increase 10/300 GL column (Cytiva). The peak fraction containing P2X3:peptidisc was analyzed by SDS-PAGE. Subsequently, the P2X3:peptidisc complex was incubated with camlipixant (Pharmaron) at a molar ratio of 1:5 for 30 min on ice before preparing cryo-EM grids.

### Cryo-EM grid Preparation and Data Acquisition

A monodisperse fraction of P2X3 was used for grid preparation. Holey carbon gold Quantifoil grids ( R1.2/1.3 mm, hole space 300 mesh) were glow-discharged for 60 s at 25 mA, holding for an additional 10 s. Subsequently, 3.0 µL of the sample was applied to the grids and blotted for 2 s at 4 °C and 100% humidity, followed by vitrification by plunging into liquid ethane cooled by liquid nitrogen using a Vitrobot Mark IV (Thermo Fisher). Micrographs were captured using a Titan Krios (Thermo Fisher) operating at 300 kV with a nominal magnification of 105,000× for all the datasets. Micrographs were obtained with a nominal defocus values of 0.8 −2.0 µm. A calibrated pixel size of 0.4125 Å was used for the processing. Videos were recorded using Leginon 3.6 (Suloway *et al*, 2005)at a dose rate of 28.56 e−/Å2/s with a total exposure time of 1.80 seconds, accumulating a dose of 56.8 e−/Å2. Intermediate frames were captured every 0.03 seconds, totaling 60 frames per micrograph. Micrographs were recorded using an automated acquisition program (EPU).

### Electron Microscopy Data Processing

Image processing and structure determination involve several steps. First, motion correction and summation of images were performed using MotionCor2 (Zheng *et al*, 2017). Contrast transfer function parameters for each non-dose-weighted micrograph were estimated as defocus values using Gctf (Zhang, 2016). Particle picking was performed using the DoG Picker (Voss *et al*, 2009), followed by initial reference-free 2D classification using CryoSPARC v.4.2.0 (Punjani *et al*, 2017). Representative 2D class averages were selected and further particle cleanup was performed with multiple rounds of 2D classification. Subsequently, initial *ab initio* reconstruction was performed using CryoSPARC. The particles were then classified into five classes using 3D classification, with the initial reconstruction low-pass filtered to 20 Å as the reference model. Multiple rounds of 3D classification were performed, and the best subset, showing clear structural features, underwent heterogeneous refinement for six rounds, resulting in a high-quality subset.

These refined particles were further subjected to nonuniform refinement (Punjani *et al*, 2020), which generated a map with the indicated global resolution at a Fourier shell correlation of 0.143. The density map was refined by local refinement with a tight mask covering only the protein region, whereas the surrounding peptidisc electron densities were manually removed using UCSF Chimera (v.1.16) (Pettersen *et al*, 2004). The final volume map was calculated with C3 symmetry using CryoSPARC, and the local resolution was estimated using Phenix (Afonine *et al*, 2018) and displayed using PyMol (DeLano, W.L.) and Chimera X (UCSF Chimera X). The final resolutions are reported in Table S1, and a detailed data-processing pipeline is shown in Supplementary Figures S3 and S8.

### Model building and structure determination

The initial apo state model of P2X3 was constructed in Coot v0.8.9 (Emsley & Cowtan, 2004) using the alphafold (Jumper *et al*, 2021) structure as a reference. The individual monomer P2X3 structures were fitted to cryo-EM electron density maps using ChimeraX, followed by successive manual adjustments and rebuilding iterations using Coot. The N-terminal domain was constructed de novo in Coot. Throughout the model-building process, manual refinements were performed based on the electron density map quality in Coot, followed by real-space refinement in PHENIX (Afonine *et al*, 2018).

In both apo and complex structures, residues ranging from 5 to 21 were missing from the models because of the absence of corresponding density in the maps. The geometric quality of the atomic models was evaluated using MolProbity (Chen *et al*, 2015). The detailed refinement statistics are provided in Table S1. The pore-lining surfaces and channels within the receptor were calculated using HOLE (Smart *et al*, 1996) and MOLE (Pravda *et al*, 2018), respectively. All visual representations were generated using UCSF Chimera or PyMOL (DeLano, W.L.).

### Structural Homology Search Using Distance Matrix Alignment (DALI)

The structure of each domain was compared with all available protein structures in the Protein Data Bank (PDB) using a heuristic PDB search on the DALI server (Holm *et al*, 2023). According to the generally accepted guidelines, a Z-score greater than 20 indicates that the two structures are “definitely homologous,” while a Z-score between 8 and 20 suggests they are “probably homologous.” Z-scores between 2 and 8 fall into a “grey area,” and those below 2 are considered not significant.

### Comparative modeling of P2X2/3 receptor binding camlipixant

Our approach involves modeling the structure of the P2X2 monomer using our camlipixant-bound P2X3 structure as a reference. Initially, five apo models of P2X2 were generated through sequence alignments obtained using Modeller10.5 (Webb & Sali, 2016), yielding the best discrete optimized protein energy (DOPE) score of −27,164. DOPE potential was derived by comparing the distance statistics from a non-redundant PDB subset of high-resolution protein structures with the distance distribution function of the reference state. Negative normalized DOPE scores of −1 or below are likely to correspond to models with the correct fold (Webb & Sali, 2021). Subsequently, these models were refined and subjected to energy minimization using Galaxy (Ko *et al*, 2012) and Molprobility (Chen *et al*, 2015), respectively.

### Preparation of Protein and Ligand Structures and Molecular Docking

For comparative analysis, three protein structures were selected: P2X7 from *Ailuropoda melanoleuca* (PDB-ID, 5U1Y), zebrafish P2X4 (PDB-ID, 8JV5), and P2X3 (in-house cryo-EM structure). Additionally, a homology model of the P2X2/3 heterotrimer (comprising of one human P2X2 monomer and two P2X3 monomers) was included. Each protein structure exists in a trimeric form with three identical ligands bound to each monomer. Specifically, each protein monomer was bound to a ligand, GW791343 (PDB-ID, 5U1Y), BX430 (PDB-ID, 8JV5), or camlipixant (in-house cryo-EM structure).

All protein structures were prepared using default parameters via the protein preparation workflow of Schrödinger Suite version 2023-4. Briefly, water molecules and other het groups, excluding ligands, were removed to maintain homogeneity in the protein-binding pockets. Hydrogen atoms were added at pH 7.4, the missing side chains were filled, and the hydrogen-bonding network was optimized using PROPKA (Olsson *et al*, 2011). Disulfide bonds were created where possible, and proteins were minimized to achieve a Root Mean Square Deviation (RMSD) gradient of 0.30 Å compared to the starting structure using the OPLS4 force field.

The prepared structures were used to generate protein grids. Active site grids were generated around ligands present in chain A of each protein structure (5U1Y, 8JV5, and cP2X3). For the P2X2/3 heterotrimer structure, an active-site grid was generated around the modeled ligand present in P2X2 (chain A) at the interface between P2X2 and P2X3 (chains B or C). The grid box center was defined as the center of mass of the respective ligand in the protein structure, with a grid box size of 27.1 Å^3^ (outer box) and 10.0 Å^3^ (inner box). The generated grids were used for molecular docking of the ligands. Additionally, the bound ligands GW791343, BX430, and camlipixant were separately extracted and saved. Ligands were prepared using the LigPrep module, where ionization states were generated at pH 7.0 ± 2.0, using Epik (Johnston *et al*, 2023), retaining input chirality.

Molecular docking calculations were performed in extra precision (XP) mode with default parameters using the Glide module of the Schrödinger Suite 2023-4. Docking of all three ligands was performed in all four structures as a matrix, and the XP scores were recorded. Subsequently, Molecular Mechanics with Generalized Born Surface Area (MM-GBSA) binding free energy was calculated for each protein-ligand complex using the Prime MMGBSA module.

### Molecular Dynamics Simulation

The docked complexes served as starting structures for molecular dynamics (MD) simulations conducted using the Desmond simulation package included in the Schrödinger Suite. The systems were built as orthorhombic solvation boxes with periodic boundary conditions in each direction, ensuring a minimum distance of 10 Å between the protein and edge of the water box. The SPC water model was employed and the systems were neutralized by adding an appropriate number of counterions.

Prior to MD simulations, Desmond’s default system relaxation protocol was used for equilibration. This included a series of stages: a 100 ps Brownian dynamics NVT simulation at 10 K with constraints on the heavy atoms of the protein, followed by 12 ps NVT simulations at 10 K and 12 ps NPT simulations at 10 K, both with restrictions on solute heavy atoms. The final relaxation stage involved increasing the temperature from 10 K to 300 K in 12 ps under NPT ensemble conditions, followed by a 24 ps NPT ensemble simulation at 300 K with no restraints on any atom. The production MD simulation length was 20 ns, with a recording interval of 100 ps, utilizing OPLS4 force-field parameters. Two complexes, namely, the P2X3-cryoEM structure complexed with camlipixant and the P2X2/3 heterotrimer homology model complexed with camlipixant at the P2X2-P2X3 interface, were simulated for 100 ns.

Upon completion of the MD simulations, trajectories were analyzed to assess ligand stability and protein conformational changes using root mean square deviation (RMSD) and protein-ligand interactions. Additionally, the change in the binding free energy (*Δ*G) was calculated using the last 5 ns of the MD simulation trajectories for each complex structure. For the two specified complexes, the MMGBSA energies were calculated based on the last 20 ns trajectory.

### Isothermal titration calorimetry (ITC) and Microscale Thermophoresis (MST) measurements

ITC measurements were conducted at 20 °C using a NANO ITC system (TA Instruments) to assess the drug binding affinity of P2X3. The compound solution was equilibrated with protein buffer D. Prior to use, all protein and compound solutions were placed in the same degassed buffer. ITC cells were loaded with 250 μl of P2X3 receptor (5 μM) and titrated against the drug compound (20 μM). Titration was performed by injecting 0.1 μl of substrate into the protein, followed by 19 injections of 0.25 μl each. The stirring rate was maintained at 200 revolutions per minute. For the control experiments, the compound was injected into the buffer alone. The equilibrium association constants were determined by fitting the reference-corrected data using the binding model provided by the manufacturer.

MST measurements were conducted at room temperature using the Monolith NT.115 system (NanoTemper Technologies). Briefly, fluorescein-labeled P2X3 was prepared in buffer D according to the manufacturer’s instructions (Red Tris 2nd generation dye, NanoTemper). A 2-fold dilution series of camlipixant was then prepared in buffer D, resulting in final concentrations ranging from 20 μM to 0.61 nM. The samples were then incubated on ice for at least 30 minutes. After incubation, samples were loaded into premium-coated capillaries (NanoTemper Technologies) and subjected to MST analysis. The results were analyzed using TJump analysis, and the obtained values were normalized and plotted against the linear concentration of camlipixant. The dissociation constant was determined by fitting the curve to a binding model provided by the manufacturer.

### Intracellular calcium influx assay

The assay was performed using the Fluo-8 Calcium Flux Assay Kit (Abcam, Cat# ab112129), following the manufacturer’s instructions. Cells were centrifuged from the culture medium and resuspended in an equal amount of Fluo-8 dye-loading solution along with Hank Balanced Salt Solution (HBSS buffer) supplemented with calcium (Gibco). A suspension containing 200,000–250,000 P2X3-Expi293 GnTI- cells per well was prepared in a 96-well poly-D-lysine plate. The dye-loading plate was then incubated in a cell incubator for 30 min at 37 °C with 8% CO_2_. ATP:Mg^2+^ was used in the range of 0 – 2 μM and camlipixant was used in the range of 0 – 2 μM in the assay. These compounds were dispensed directly onto cell plates and the data were collected simultaneously. The maximum signal was used to generate a plot. A fluorescence-based assay was used to detect intracellular calcium mobilization in cells at excitation/emission wavelengths of 490/525 nm using a microplate reader G5 machine (SpectraMax). The readings obtained from the blank standard were used as negative controls. This value was subtracted from the readings of other standards to obtain baseline-corrected values.

## Supporting information

Combined-supplementary-data-file

## DATA AND CODE AVAILABILITY

Three-dimensional cryo-EM density maps and coordinates for camlipixant-bound P2X3 (accession codes EMD-44771 and 9BPC) and ATP-bound P2X3 (accession codes EMD-44772 and 9BPD) were deposited in the Protein Data Bank. The overall map was used to refine the ATP-bound P2X3 structure. Additionally, focused refinements were conducted on both the extracellular domain (excluding the transmembrane domain) and transmembrane domain (excluding the extracellular domain) to enhance the visualization of specific features present in the overall map. Although these refined maps aided in model building, they were not utilized for refinement in PHENIX.

## ACKNOWLEDGEMENT

Special thanks go to Dr. Robert V. Stahelin for granting access to the ITC machine, Dr. T. Klose and Dr. F. Vago for cryo-EM and S. Wilson for computation. We acknowledge the Purdue cryo-EM facility for access to instrumentation for data collection. This work was partly supported by the Indiana Clinical and Translational Sciences Institute, funded by Grant Number UM1TR004402 from the National Institutes of Health, National Center for Advancing Translational Sciences, and Clinical and Translational Sciences Award to Thach T.

## Author Contributions

TT and KD conducted the cellular, biophysical, and structural studies. PPN performed all the MD simulations. SS contributed to the creation of the stable cell line. TT, IH, SA, and JS were involved in designing the experiments, specifically the selection of inhibitors and the study design. JS and RS developed this project. RS was involved in the structure determination, map interpretation, and result analysis. All authors participated in manuscript preparation and review.

