## Supplementary material for "A Second Drug Binding Site in P2X3": Combined-supplementary-data-file

**CONTENTS**

1. SUPPLEMENTARY TaBLES

Table S1. Cryo-EM data collection, refinement, validation and statistics of cP2X3

Table S2. Residues in the Camlipixant binding pocket and sequence alignment across the P2X receptor family and its conservation.

Table S3. Molecular dynamic simulations among P2X members with different ligands.

Supplementary Figures

Figure S1. eGFP-P2X3 expression in the Expi293F GnTI^-^cells.

Figure S2. Purification of P2X3.

Figure S3. The cryo-EM data processing pipeline for P2X3:ATP reconstituted into DDM.

Figure S4. The cryo-EM structure of ATP-bound cP2X3 was superimposed with the crystal structure of hP2X3:ATP.

Figure S5. The binding of camlipixant to P2X3.

Figure S6. The binding affinity for ATP towards P2X3.

Figure S7. The impact of peptidisc in P2X3 structure determination.

Figure S8. The cryo-EM data processing pipeline for P2X3:camlipixant reconstituted into peptidisc.

Figure S9. Cryo-EM density maps for the camlipixant-bound P2X3 structure.

Figure S10. Conformational changes in camlipixant-bound P2X3 structure.

Figure S11. Sequence alignment of *Canine lupus* P2X3 (cP2X3) and Human P2X3 (hP2X3).

Figure S12. The sequence alignment of cP2X3 and hP2X2 isoforms.

Figure S13. The drug-binding pocket enlarges in the Cam-bound P2X3 receptor.

Figure S14. Molecular dynamic simulations of camlipixant-bound P2X2/3 receptor.

Figure S15. A model of the P2X2/3-gefapixant complex.

Figure S16. The Cam binding site appears across P2X receptors.

Figure S17. Molecular dynamics (MD) simulations of camlipixant with P2X7, P2X4, and our camlipixant-bound P2X3.

Table S1. Cryo-EM data collection, refinement, validation and statistics.

| **Structures** | **P2X3:ATP in DDM**  PDB-ID, 9BPD  EMD-44772 | **P2X3:Cam in DDM** | **P2X3:Cam in peptidisc**  PDB-ID, 9BPC  EMD-44771 |
| --- | --- | --- | --- |
| **Data collection andprocessing** |  | | |
| Magnification | 105,000 | 105,000 | 105,000 |
| Voltage (kV) | 300 | 300 | 300 |
| Electron exposure ((e–/Å2) | 56.8 | 56.8 | 56.8 |
| Defocus range ((μm) | 0.8-2.0 | 0.8-2.0 | 0.8-2.0 |
| Raw pixel size ( Å) | 0.411 | 0.411 | 0.411 |
| Symmetry imposed | C3 | C3 | C3 |
| Number of initial particle images | 87,363 | 561,446 | 200,982 |
| Number of final particle images | 44,237 | 373,772 | 40,142 |
| Map resolution (Å) | 3.63 | 2.93 | 3.44 |
| FSC threshold | 0.143 | 0.143 | 0.143 |
| Map resolution range ( Å) | 3.0-5.0 | 2.5-5.0 | 3.0-5.0 |
| **Refinement** |  |  |  |
| Initial model used | Alphafold model | Apo model | Apo model |
| Model resolution ( Å) | N/A | 3.6 | 3.6 |
| *FSC threshold* | 0.5 | 0.5 | 0.5 |
| *Model resolution range (* Å) | N/A | 3.0-50 | 3.0-50 |
| *Map sharpening B factor (* Å^2^) | -100 |  | -90 |
| *Model composition* |  |  |  |
| *Non-hydrogen atoms* | 7551 |  | 7362 |
| *Protein residues* | 918 |  | 927 |
| *Ligand: Mg* | 3 |  |  |
| *ATP* | 3 |  |  |
| *Camlipixant* |  |  | 3 |
| NAG | 15 |  | 9 |
| B factors *(*Å^2^) |  |  |  |
| Protein | 160.45 |  | 85.50 |
| Ligand | 168.09 |  | 80.68 |
| RMSD values |  |  |  |
| Bond lengths ( Å) | 0.003 |  | 0.004 |
| Bond angles (^o^) | 0.634 |  | 0.612 |
| Validation |  |  |  |
| Molprobity score | 2.63 |  | 2.09 |
| Clash score | 13.18 |  | 8.51 |
| Poor rotamer (%) | 0.38 |  | 1.20 |
| Ramachandran plot (%) |  |  |  |
| Favored | 87.50 |  | 89.04 |
| Allowed | 12.17 |  | 10.96 |
| Outliers | 0.33 |  | 0 |

| cP2X3 | HUMAN | | | | | |
| --- | --- | --- | --- | --- | --- | --- |
|  | P2X3 | P2X2 | P2X4 | P2X5 | P2X6 | P2X7 |
| Y65 | **Y70** | L/S | T75 | T76 | T59 | V64 |
| R68 | **R73** | H/K | **R83** | **R84** | **R65** | S78 |
| M70 | **M75** | G/W | W85 | W86 | W67 | F80 |
| I88 | **I93** | I/- | V102 | V104 | L82 | V106 |
| M91 | **M96** | V/- | V105 | L107 | F106 | F111 |
| M160 | **M165** | G | F179 | L182 | L137 | L184 |
| F277 | **F282** | F | **F297** | **F299** | T245 | Y295 |
| Y280 | **Y285** | Y | **Y300** | **Y302** | H248 | **Y305** |
| E288 | **E293** | T | **E308** | **E310** | **E256** | **E313** |
| L293 | **L298** | I | I313 | M315 | **L261** | I318 |

Table S2. Residues in the Camlipixant binding pocket and sequence alignment across the P2X receptor family and its conservation.

There are ten residues involved in the camlipixant binding pocket (left column) in cP2X3. The conserved residues of the same ten residues were highlighted in different human P2X proteins (right columns). The highly conserved residues are in Bold.

Table S3. **Molecular dynamic simulations among P2X members with different ligands.**

Molecular dynamic simulations extracted values of MMGBSA Binding free energy (kcal/mol) shown as heatmap. The light color (white) represents the lowest energy, and the dark color (dark red) represents the highest energy.

| **Protein**  **Ligand** | **P2X7-5U1Y** | **P2X4-8JV5** | **P2X3** | **P2X3-P2X2 Interface** | **P2X2-P2X3 Interface** |
| --- | --- | --- | --- | --- | --- |
| GW791343 | -67.38 ± 6.04 | -55.36 ± 3.98 | -50.72 ± 3.95 | -52.57 ± 3.83 | -55.27 ± 3.95 |
| BX430 | -56.49 ± 5.99 | -58.54 ± 2.99 | -36.69 ± 4.91 | -54.66 ± 4.61 | -58.69 ± 6.08 |
| Camlipixant | -61.92 ± 3.72 | -72.54 ± 4.32 | -74.53 ± 3.32 | -68.25 ± 4.64 | -47.58 ± 4.69 |

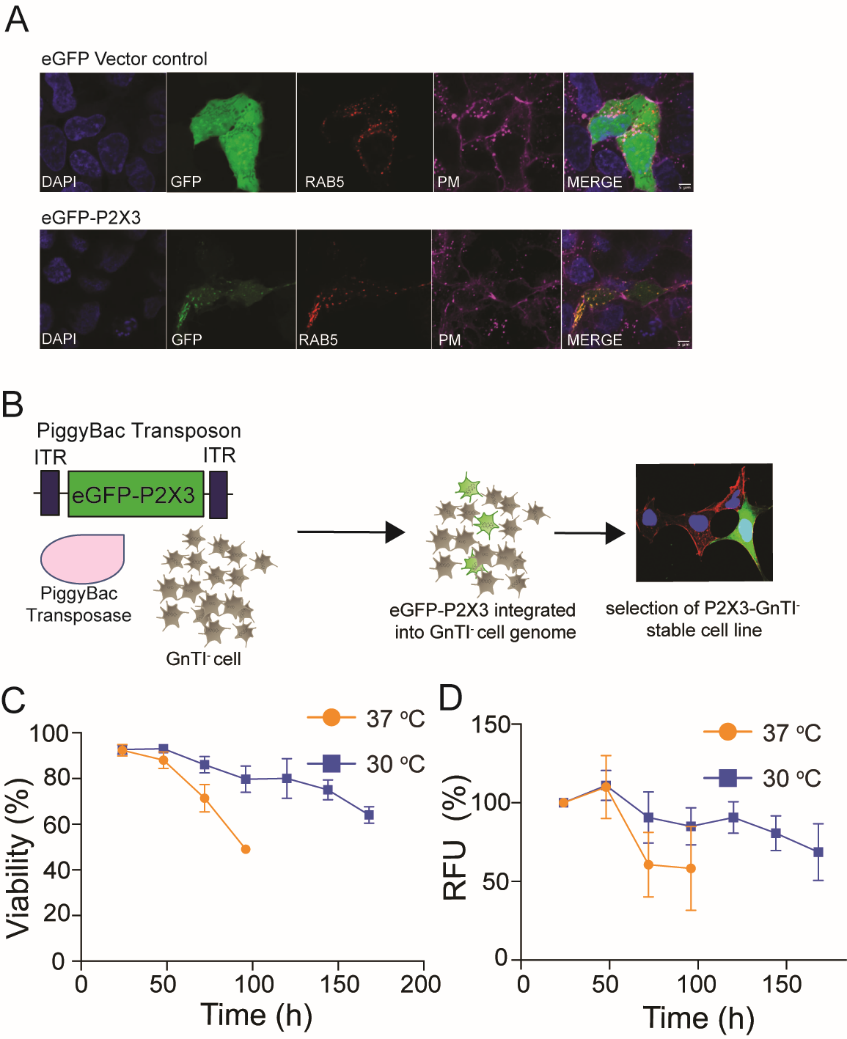

Figure S1. eGFP-P2X3 expression in the Expi293F GnTI-cells. (A) eGFP expression was monitored using fluorescence confocal microscopy. Endosomes and the plasma membrane are identified through immunostaining using a Rab5 antibody and phalloidin staining, respectively. (B) The generation process of the Expi293F GnTI-stable pool through the piggyBac transposon system is outlined. ITR, inverted terminal repeat sequence. (C) The viability and relative fluorescence unit (RFU) of eGFP-P2X3 expression are depicted at 37 °C and 30 °C.

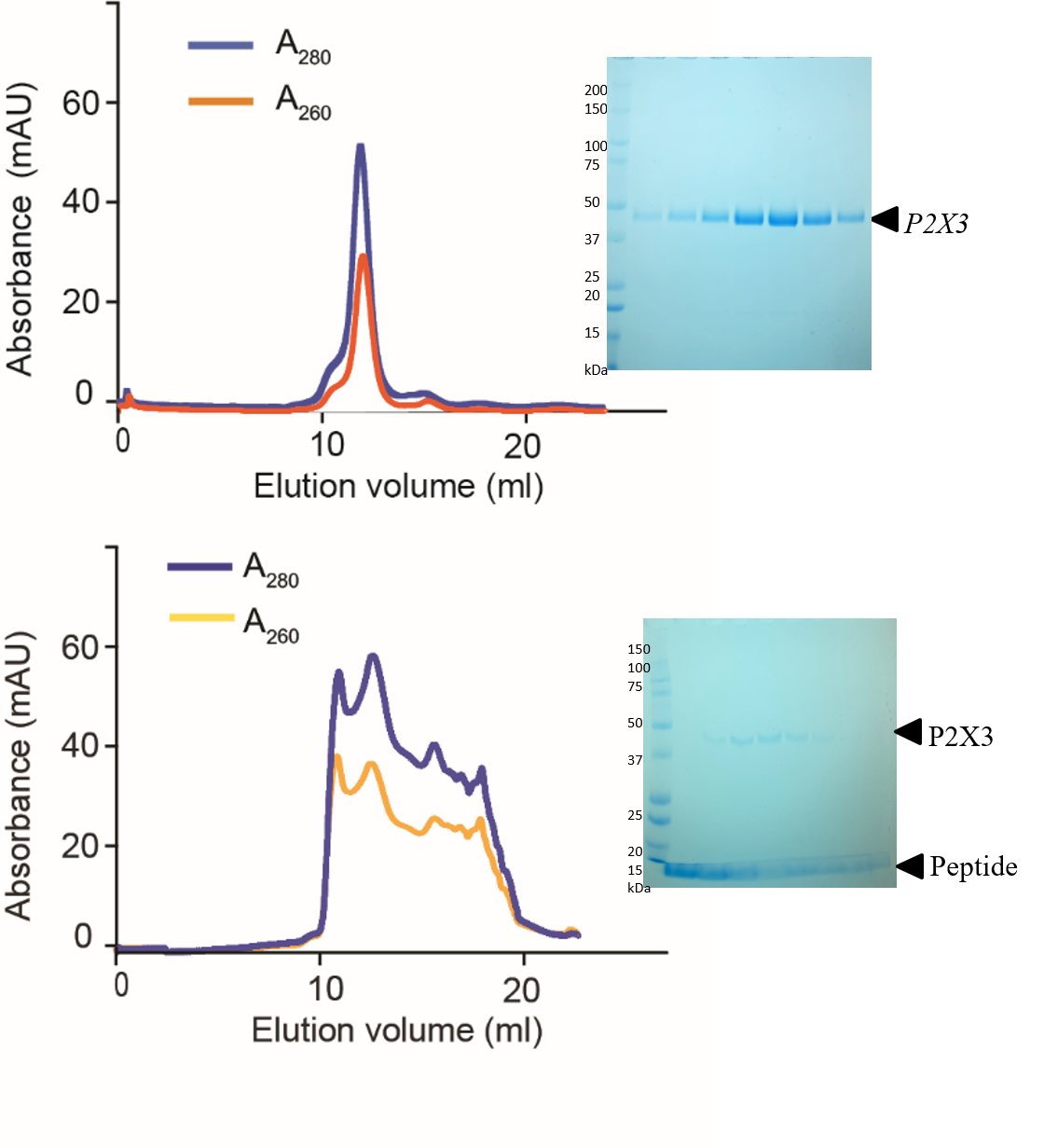

Figure S2. Purification of P2X3. (A) SEC profile of purified P2X3 and (B) SEC profile of P2X3:peptidisc complexes. Protein samples were fractionated using a Superdex 200 10/300 column (left), while the purity of the samples was assessed by SDS-PAGE (right).

Figure S2. Purification of P2X3. (A) SEC profile of purified P2X3 and (B) SEC profile of P2X3:peptidisc complexes. Protein samples were fractionated using a Superdex 200 10/300 column (left), while the purity of the samples was assessed by SDS-PAGE (right).

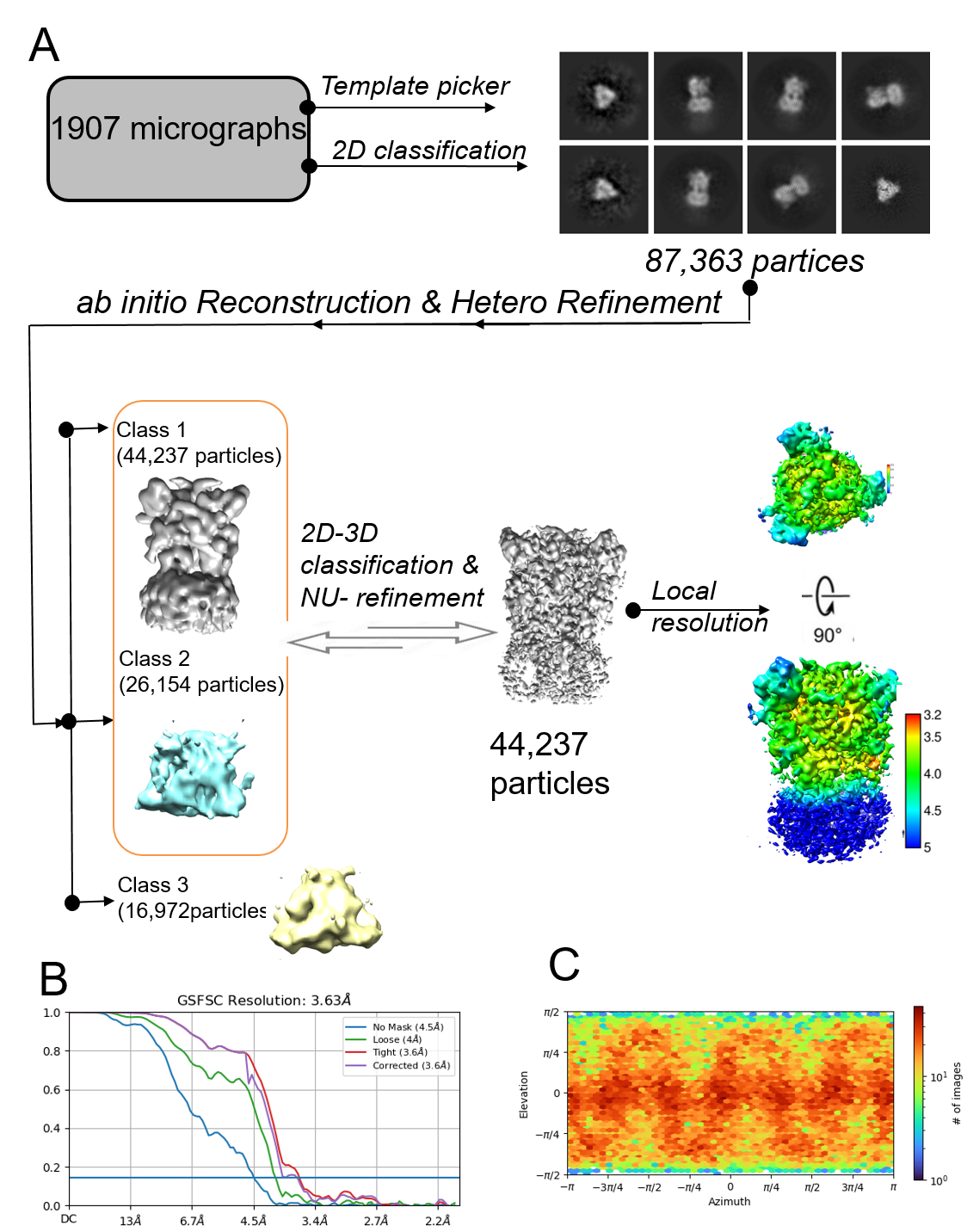

Figure S3. The cryo-EM data processing pipeline for P2X3:ATP reconstituted into DDM. (A) An overview of the processing steps that resulted in a density map with an overall resolution of 3.63 Å. A local resolution map, ranging from 3.2 to 5 Å resolution, was calculated with cryoSPARC. This map reveals that the lowest resolution (depicted in dark blue) corresponds to the transmembrane helices, as observed in both side and top views from the extracellular side. The maps were generated using UCSF ChimeraX. (B) Gold-standard Fourier shell correlation (FSC) curves were generated for resolution estimation. (C) Heat map showing the angular distribution of the particles used for the final map.

Figure S4. The cryo-EM structure of ATP-bound cP2X3 was superimposed with the crystal structure of hP2X3:ATP. (highlighted in pink, PDB-ID, 5VSK). Cartoon representations illustrate both structures, revealing a high similarity with minimal conformational changes observed in the ATP-binding site within the upper body domain. Each subunit of the cryo-EM structure of ATP-bound P2X3 is color-coded as depicted in Figure 3. (A) side view, (B) top view.

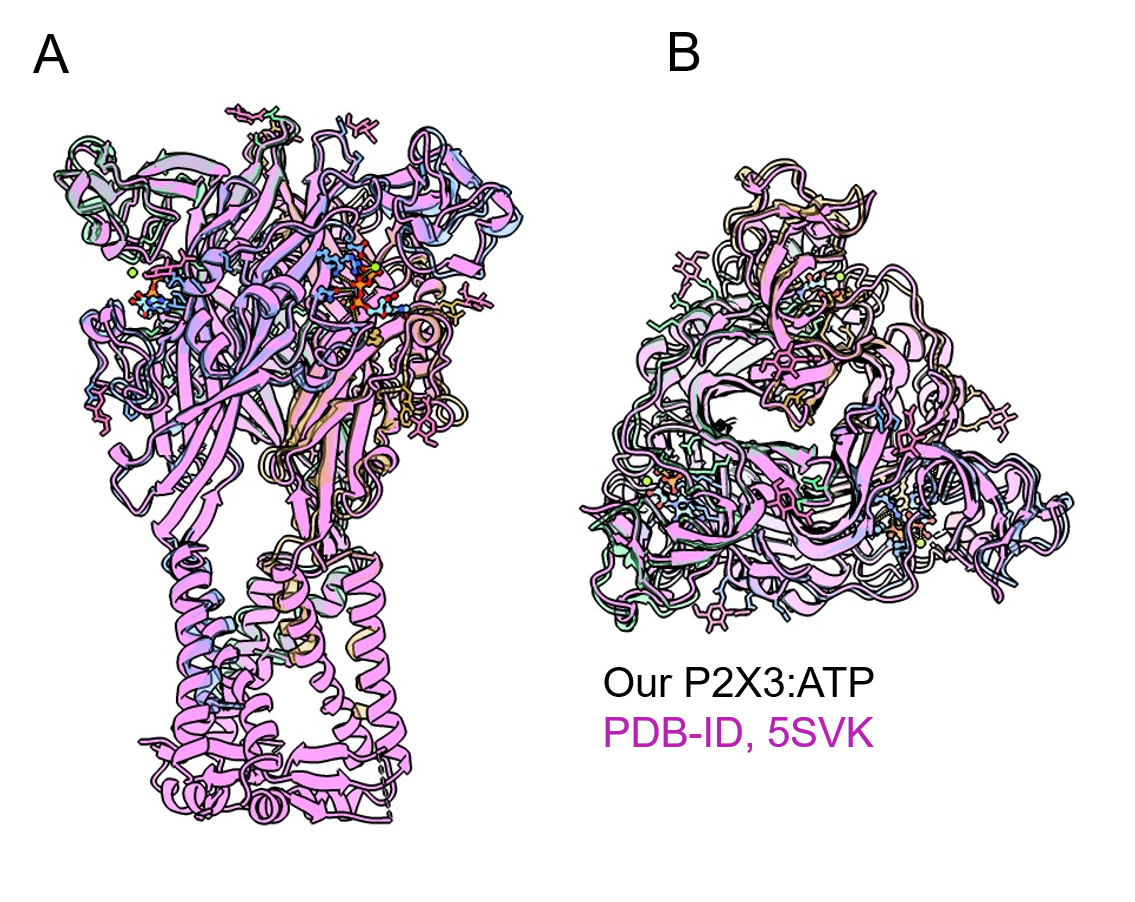

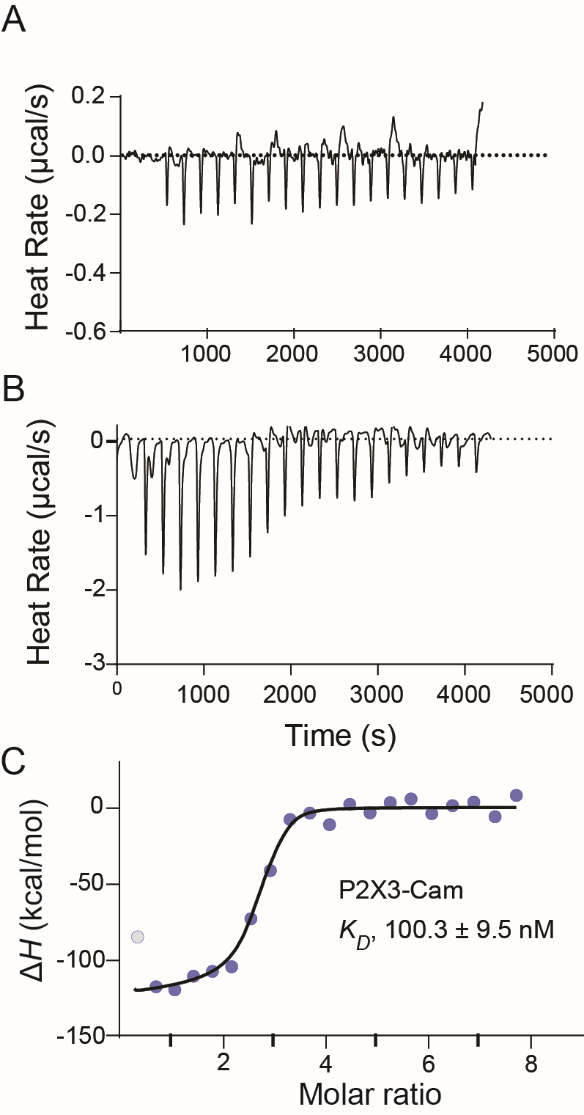

Figure S5. The binding of camlipixant to P2X3 was determined by ITC ITC curves for titrations of Cam into either P2X3:ATP (A) or P2X3 apo (B), and integrated binding isotherms are shown (C).

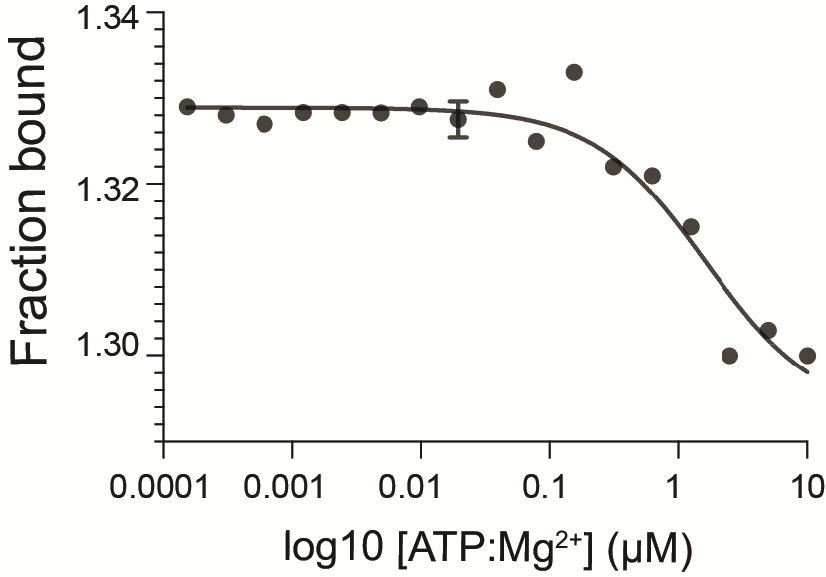

Figure S6. The binding affinity for ATP towards P2X3 was determined using MST.Integrated binding thermophoresis is displayed for the interaction. A representative binding thermophoresis displayed to show the binding nature of ATP toward cP2X3.

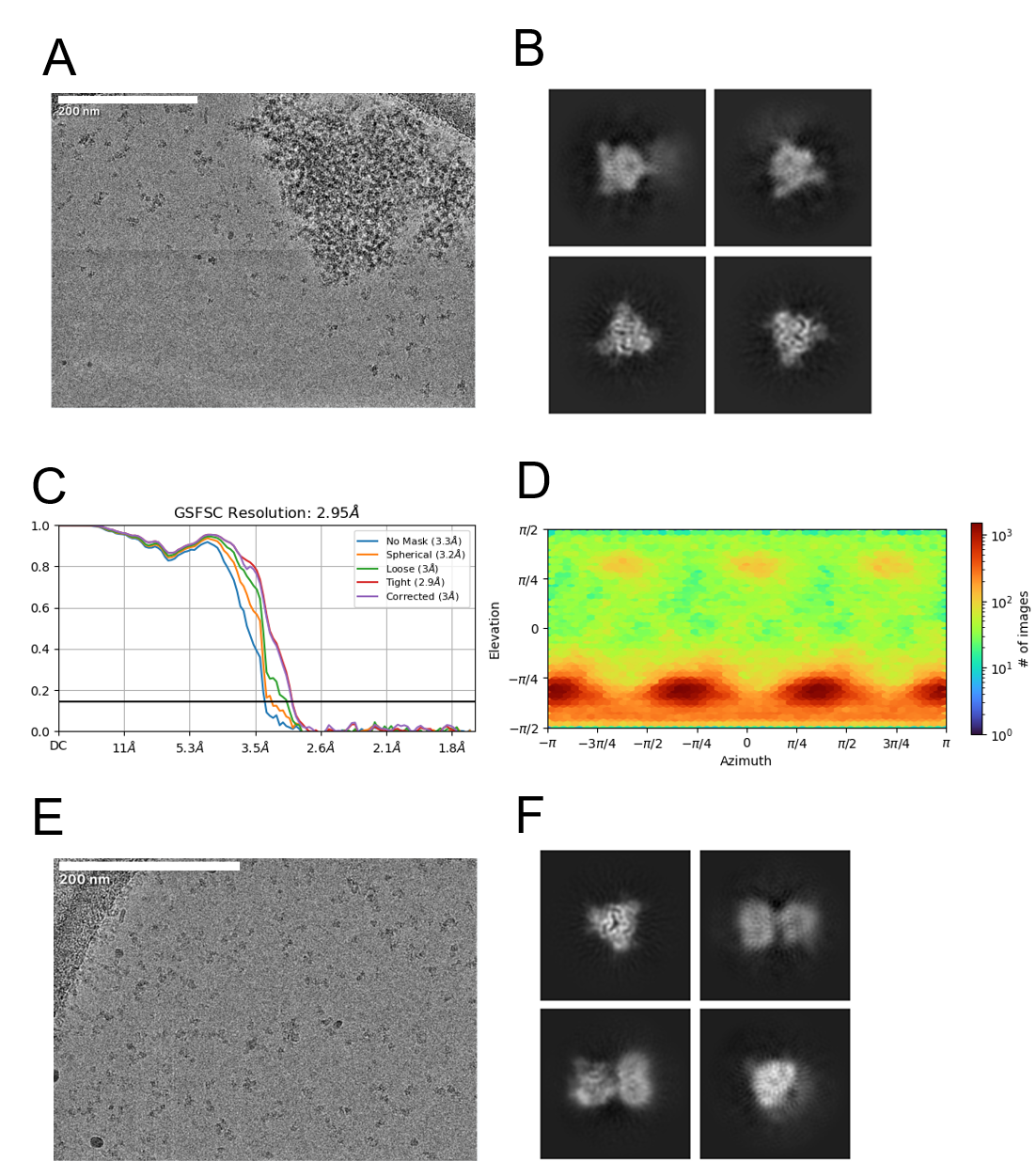

Figure S7. The impact of peptidisc in P2X3 structure determination (A) Representative cryo-EM image of P2X3:camlipixant particles reconstituted into DDM. (B) 2D classification of P2X3:camlipixant reconstituted into DDM. The gold-standard Fourier shell correlation (FSC) curves for resolution estimation are displayed in (C), while the angular distribution of the particles used for the final map is illustrated in (D). (E) Representative cryo-EM image of P2X3:camlipixant particles reconstituted into peptidisc. (F) 2D classification of P2X3:camlipixant reconstituted into peptidisc is depicted.

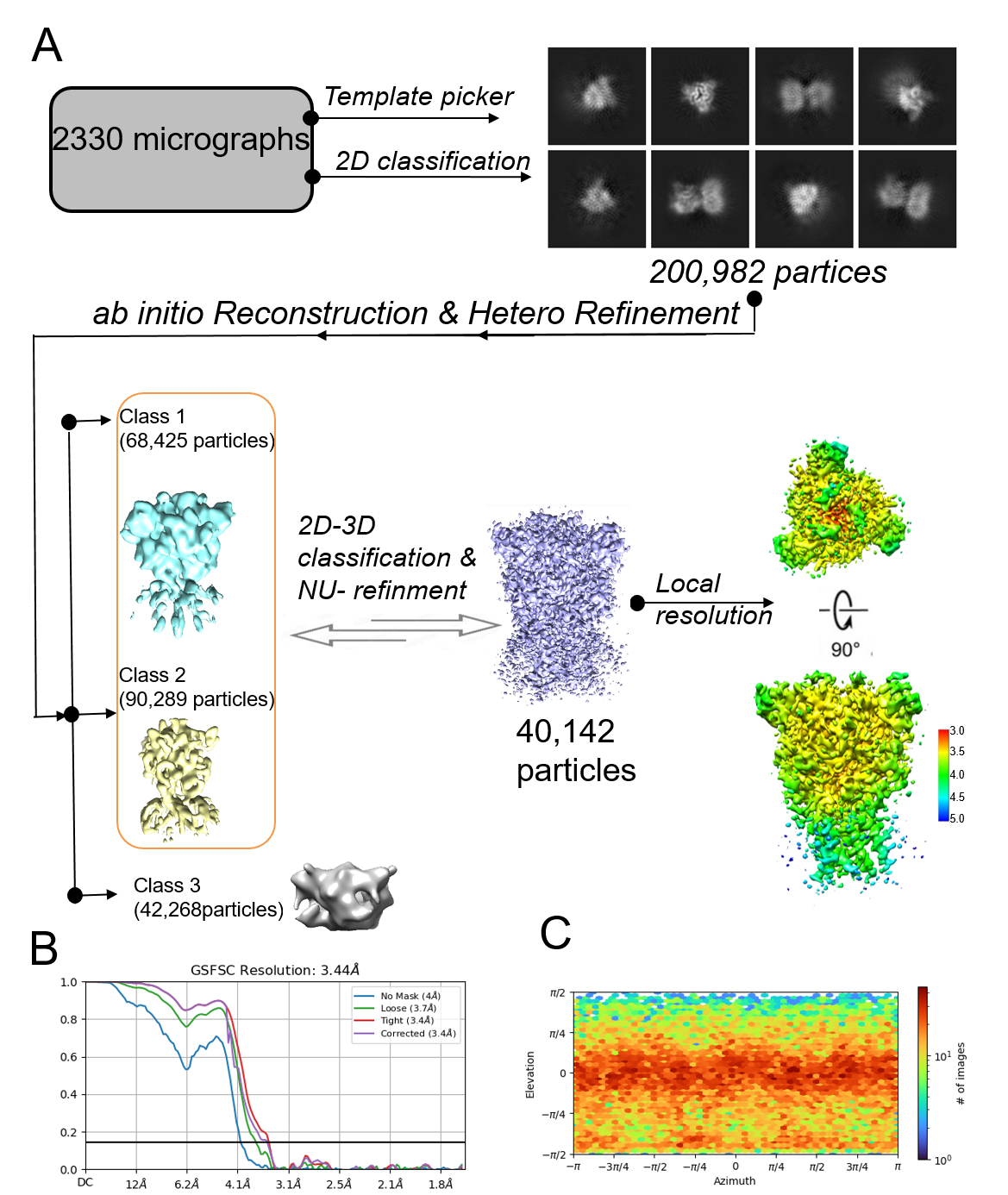

Figure S8. The cryo-EM data processing pipeline for P2X3:camlipixant reconstituted into peptidisc . An overview of the processing steps led to the generation of an electron density map with an overall resolution of 3.44 Å, utilizing 20% of the initially exported particles. Additionally, a local resolution map ranging from 3.2 to 5 Å resolution was calculated with cryoSPARC. This map indicates that the lowest resolution (depicted in dark blue) correlates with the transmembrane helices, as observed in both side and top views from the extracellular side. The maps were generated using UCSF ChimeraX. (B) Gold-standard Fourier shell correlation (FSC) curves were generated for resolution estimation. (C) The angular distribution of the particles used for the final map is illustrated.

Figure S9. Electron density maps for the camlipixant-bound P2X3 structure. (A) Overall P2X3 structure and transmembrane helixes (B). The maps depict the structure with contours set at 3.0 σ, with a carve of 2 Å. Residues within the transmembrane helix region are visualized as sticks and spheres.

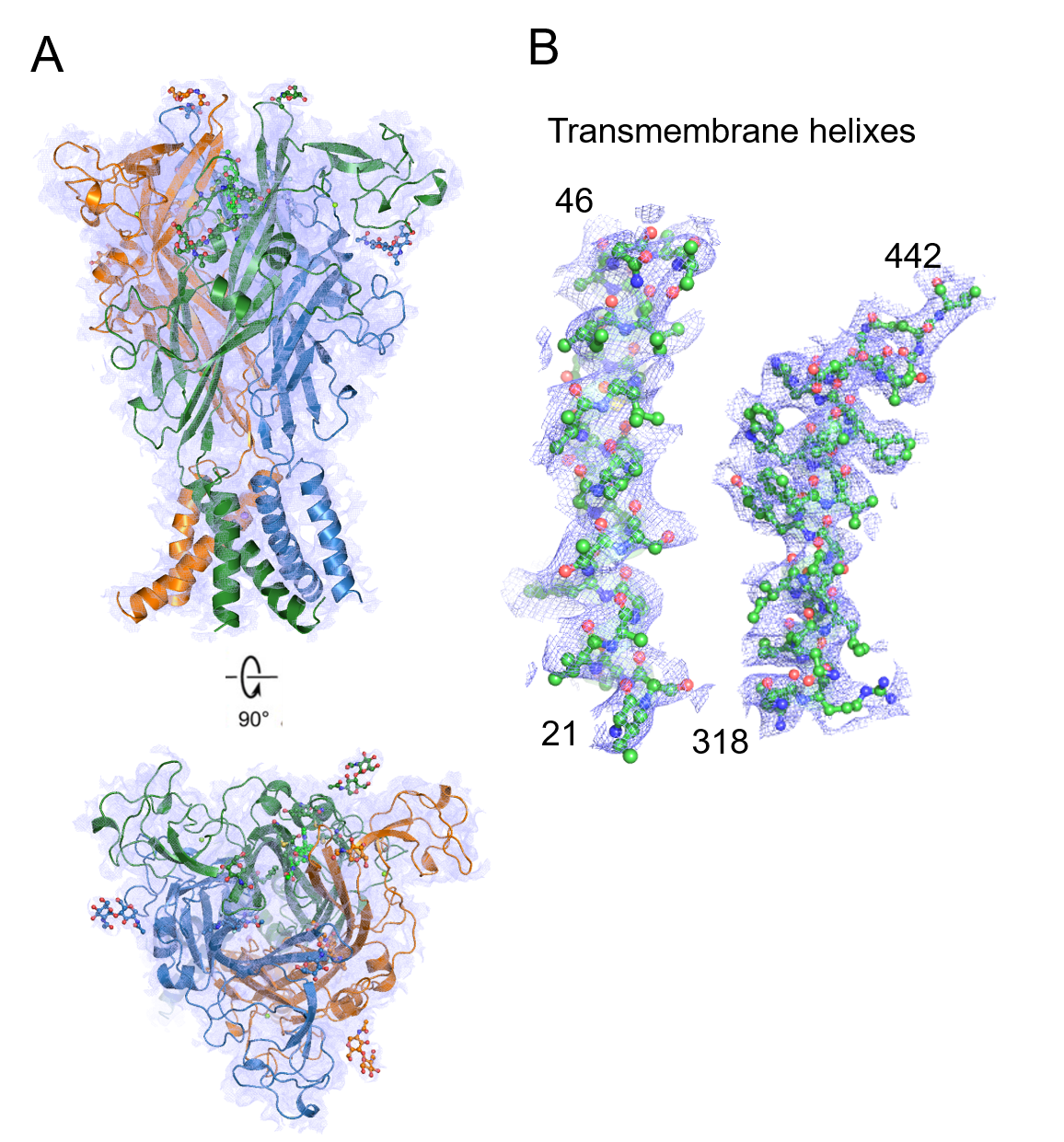

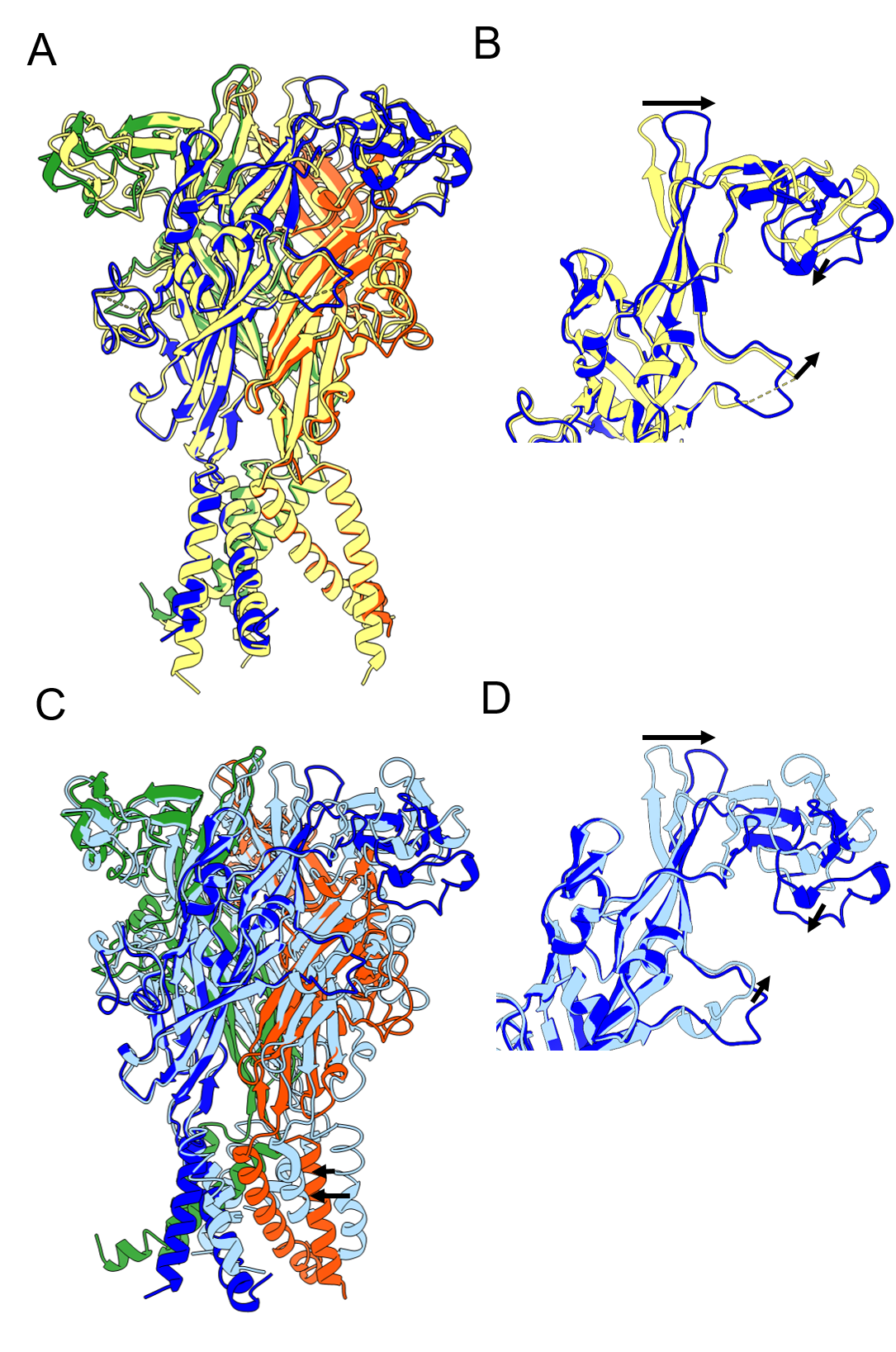

Figure S10. Conformation changes in camlipixant-bound P2X3 structure. (A, B) A comparison is made between the P2X3 receptor reconstituted into peptidisc and the P2X3 apo reconstituted into DDM (depicted in yellow, PDB-ID, 5SVJ). The structures, represented as cartoons, were superposed using ChimeraX. The overall structures appear similar, particularly in the transmembrane helices, indicating that reconstituting Cam-bound P2X3 into peptidisc preserves its structure. However, there is a conformational change upon camlipixant binding, evidenced by the enlargement of the Cam binding site and the narrowing down of the ATP binding site. (C, D) A comparison is made between the P2X3 receptor in the Cam-bound and ATP-bound states (highlighted in light cyan). Upon Cam binding, there is a conformational change characterized by the enlargement of the Cam binding site and the narrowing down of the ATP binding site, while the transmembrane helices move closer. The black arrows highlight the direction of the domain movement upon ATP binding.

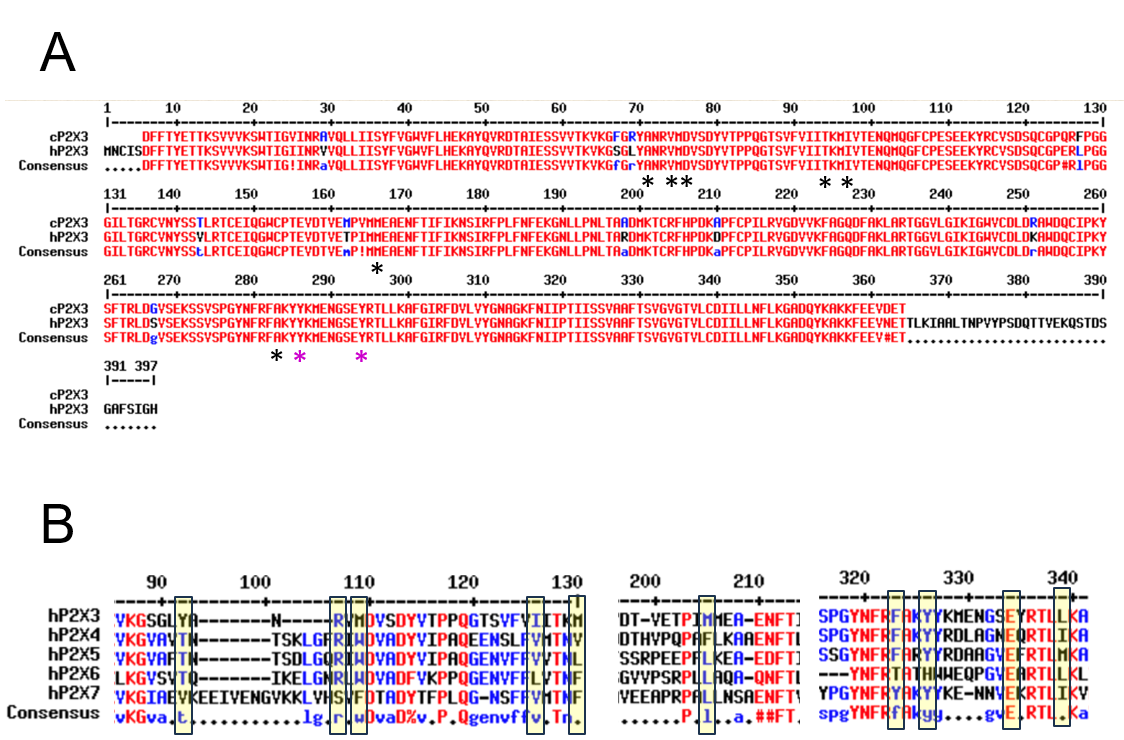

Figure S11. Sequence alignment of *Canine lupus* P2X3 (cP2X3) and Human P2X3 (hP2X3).(A). Black and pink stars indicate key residues in different subunits binding to camlipixant. Panel (B) shows the sequence alignment of hP2X3-7 receptors. Residues involved in the Cam binding pocket are highlighted within a yellow box. hP2X3 (NP_002550.2), hP2X4 (NP_001243725.1), hP2X5 (NP_001412012.1), hP2X6 (NP_001381624.1), hP2X7 (NP_002553.3).

Figure S12. The sequence alignment of cP2X3 and hP2X2 isoforms. Residues binding to camlipixant are highlighted within yellow boxes, while residues binding to gefapixant are highlighted within cyan boxes. hP2X2-I (NP_036358.2), hP2X2-H (NP_777361.1), hP2X2-J (NP_001269093.1), hP2X2-K (NP_001269094.1).

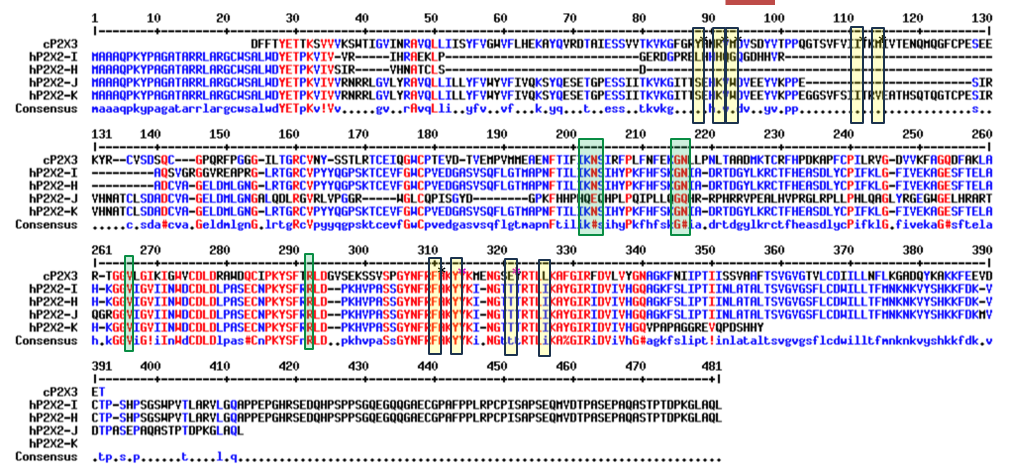

Figure S13. The drug-binding pocket enlarges in the Cam-bound P2X3 receptor. (A) Line representation of the P2X3 structure, highlighting the ATP binding pocket (purple) and the Cam binding pocket (orange). The top views of P2X3:ATP (A), apo P2X3 (B), and P2X3:camlipixant (C) are displayed. (D-F) Dot representations of P2X3:ATP (D), apo P2X3 (E), and P2X3:camlipixant (F) illustrate the internal-space turret along the molecular threefold axis running through the center of the structures. These dot plots are generated using HOLE, where blue represents a radius greater than 2.3 Å, green indicates a radius between 1.15 and 2.3 Å, and red signifies a radius less than 1.0 Å.

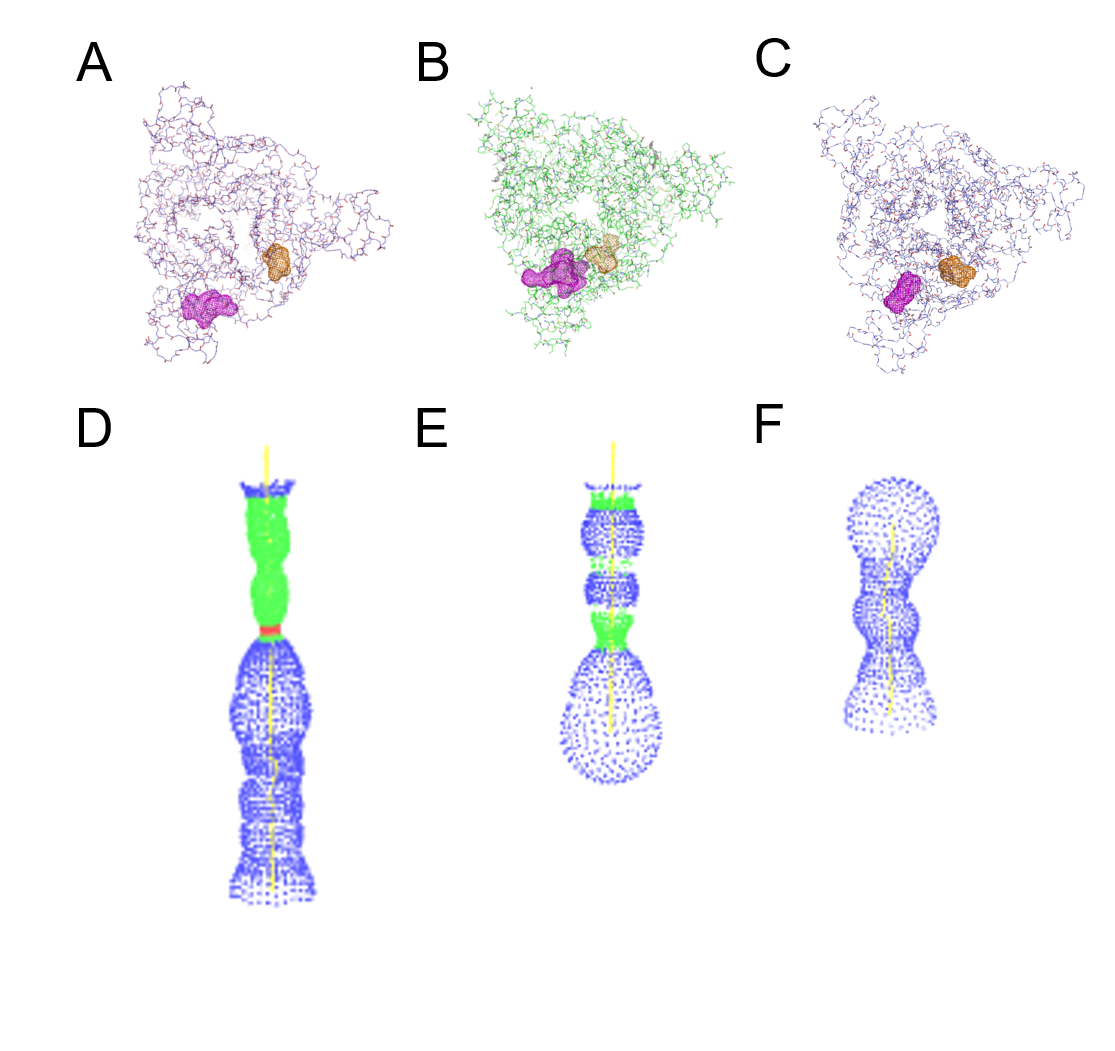

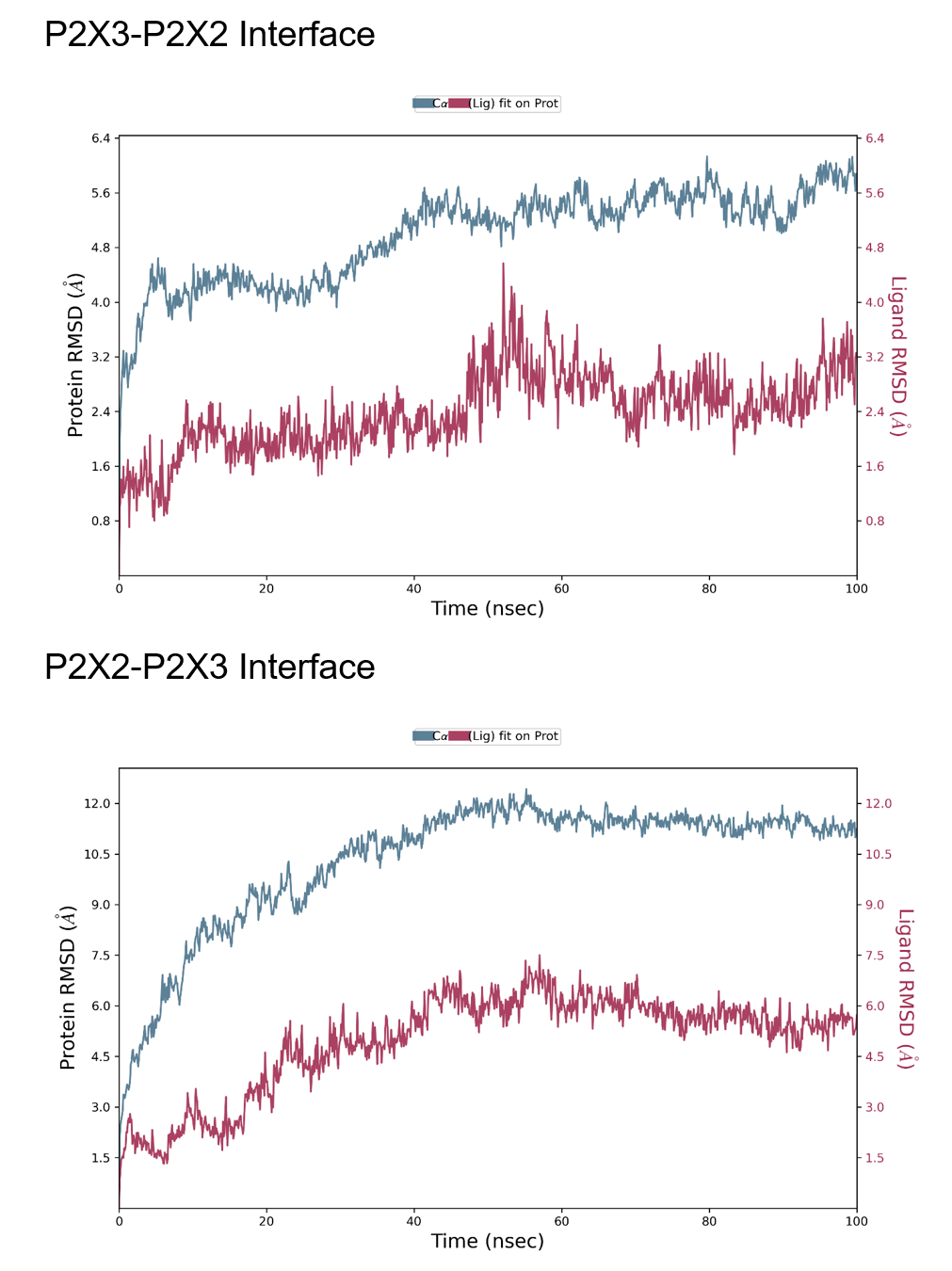

Figure S14. Molecular dynamic simulations of camlipixant-bound P2X2/3 receptor. The interactions are conducted for 100 ns. RMSD values are shown for both protein and each camlipixant ligand.

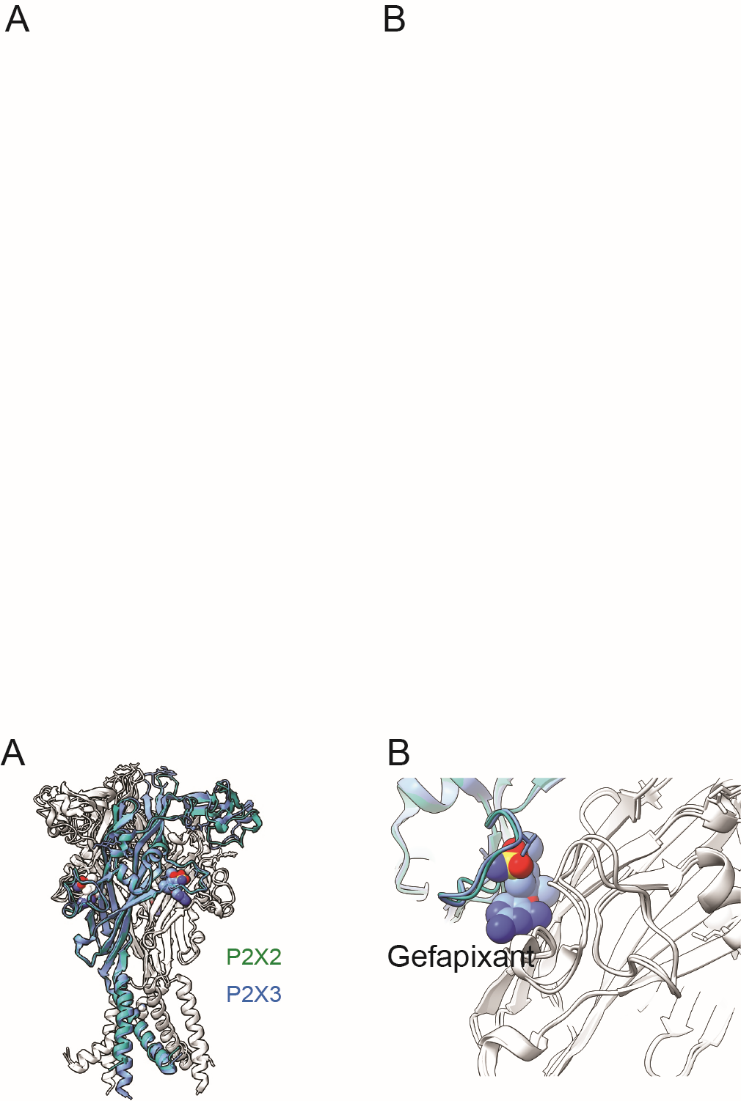

Figure S15. A model of the P2X2/3-gefapixant complex. (A)The P2X2/3 receptor superposed onto the P2X3 receptor (PDB-ID, 5YVE), represented in a cartoon diagram. (B)The gefapixant ligand is depicted as a ball-and-stick model.

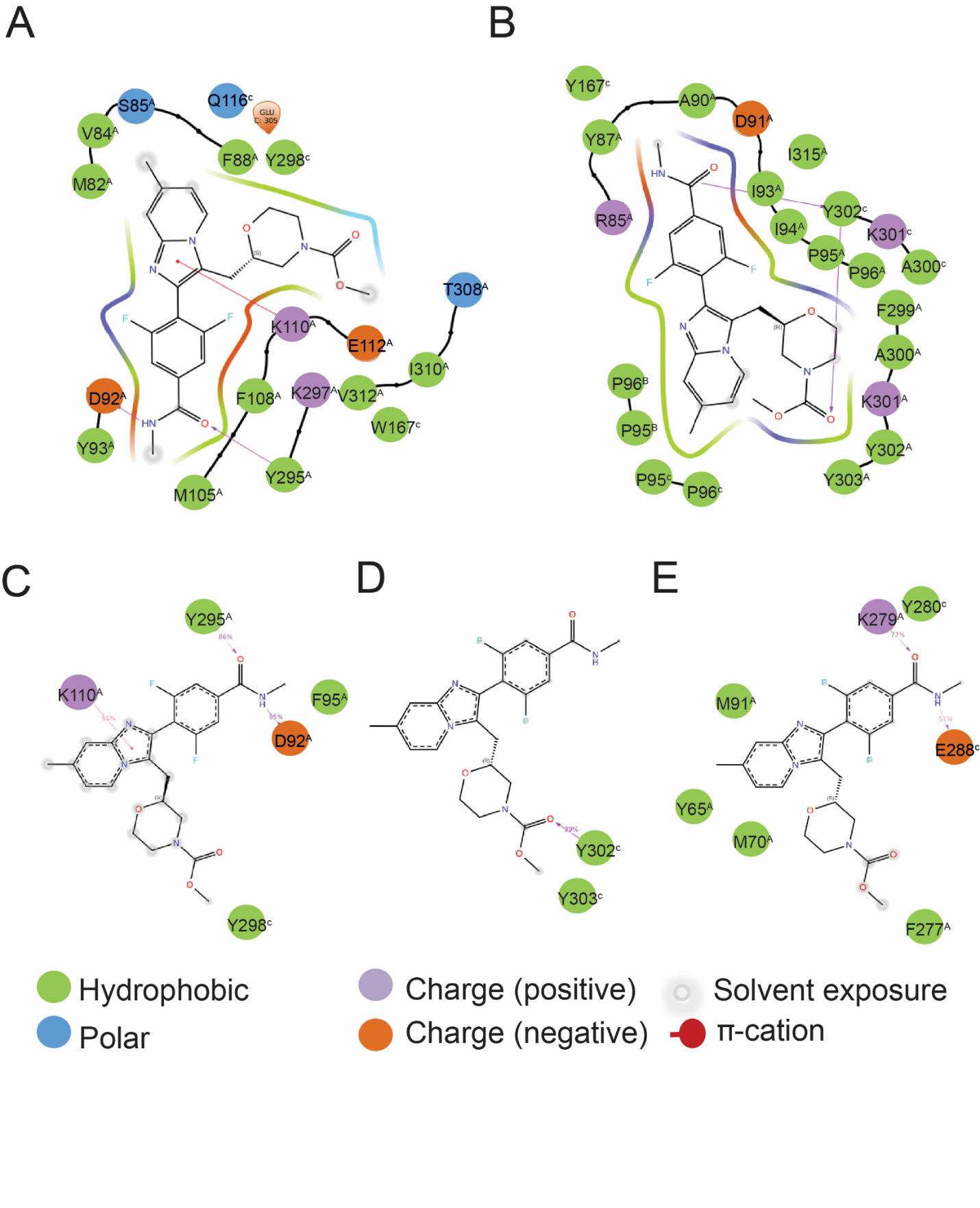

Figure S16. The Cam binding site appears across P2X receptors. Cam occupies the same binding site with a similar pose and orientation in P2X3 (Figure 3E), as observed in (A) P2X7 (PDB-ID, 5U1Y) and (B) P2X4 (PDB-ID, 8JV5). Molecular dynamic simulations of camlipixant-bound P2X receptor interactions are conducted for P2X7 (C), P2X4 (D), and P2X3 (E).

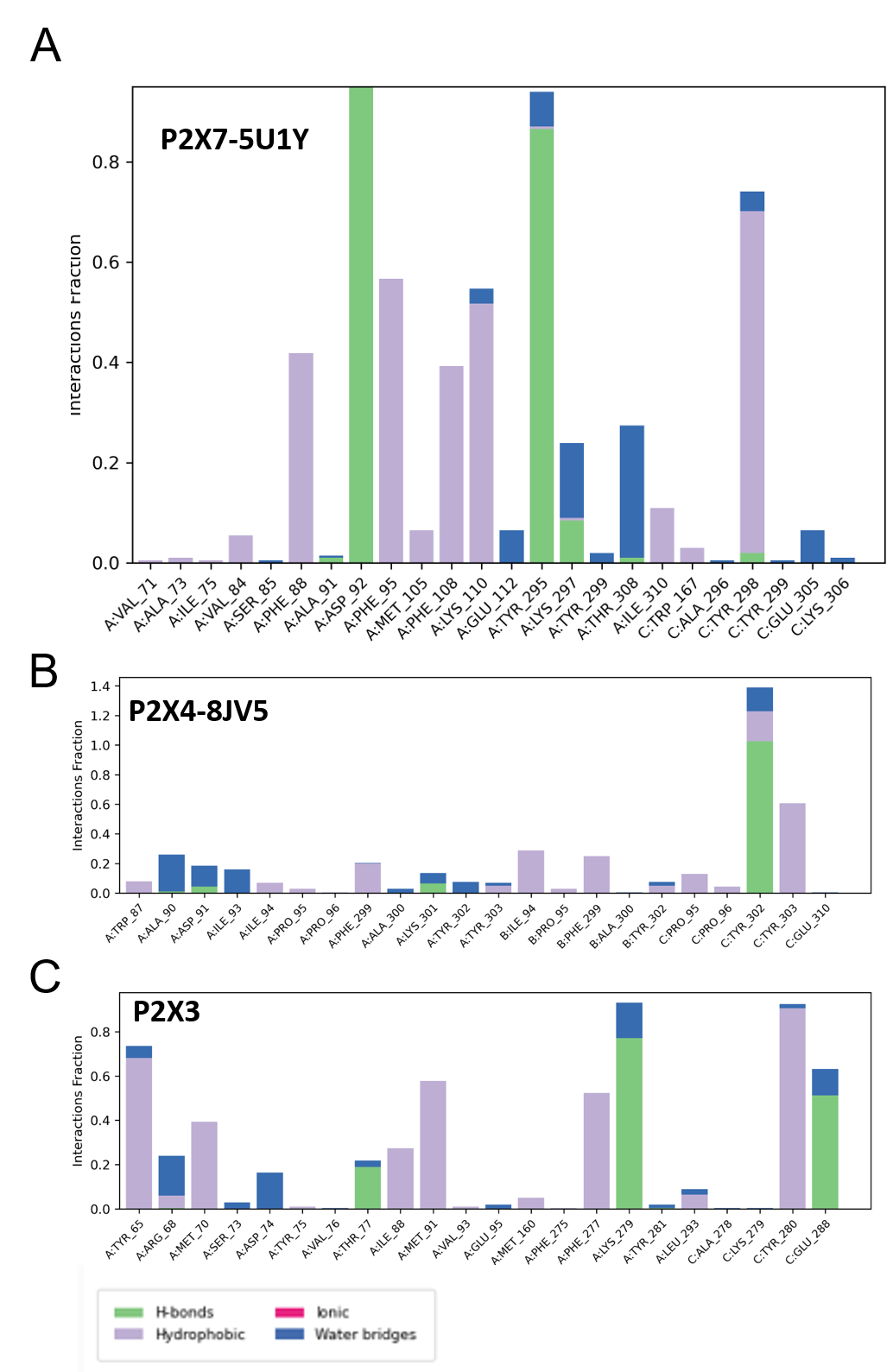

Figure S17. Molecular dynamics (MD) simulations of camlipixant with P2X7, P2X4, and our camlipixant-bound P2X3. MD simulations of camlipixant with P2X7, P2X4, and our camlipixant-bound P2X3. (A - C) with different residues in the binding pocket formed by the upper body domain reveal interactions of camlipixant with P2X7, P2X4, and our camlipixant-bound P2X3.

Video 1: Conformational changes in P2X3 when bound to camlipixant compare to P2X3 Apo structure

Video 2: Conformational changes in P2X3 when bound to camlipixant compare to ATP bound structure
